# Complex HPV-human DNA structures revealed by large-scale DNA analyses in an HPV-cancer derived cell line

**DOI:** 10.1101/2025.10.06.680684

**Authors:** Eleanor J. Agosta, Yoke-Chen Chang, Vandya Rao, Jessie Hollingsworth, Michelle Brown, Debajyoti Kabiraj, Mark Einstein, Anne Van Arsdale, Koenraad Van Doorslaer, Chang Chan, Subhajyoti De, Advaitha Madireddy, Brian J. Haas, Danny E. Miller, Jack Lenz, Cristina Montagna

## Abstract

Most human papillomavirus (HPV)-associated cancers harbor viral DNA integrated into the human genome as extrachromosomal circles, intrachromosomal segments, or both. Distinguishing intrachromosomal from identical-sequence extrachromosomal DNA (ecDNA) by sequencing alone is challenging, and the architecture of large-scale HPV-human DNA structures remains incompletely understood. To address this, we applied complementary genomic tools, spanning single-nucleotide to megabase resolution, to the HPV16-positive oropharyngeal cancer-cell line UM-SCC-47. These revealed that an initial integration event formed a 23 kb extrachromosomal heterocatemer circle comprising 7.5 kb of HPV16 DNA and 16 kb of the human *TP63* gene. Subsequent genomic rearrangements generated heterocatemer tandem arrays extending to 0.6 megabases, plus additional large-scale rearrangements involving the HPV-*TP63* structures, as revealed by long-read DNA sequencing and optical genome mapping. Fluorescent *in situ* Hybridization (FISH) showed that the heterocatemers were intrachromosomally localized at chromosome 3 at the *TP63* locus in 100% of the cells. Long-read RNA sequencing further showed that these intrachromosomal templates produced spliced, polyadenylated transcripts. A subset of cells also harbored HPV16 ecDNA derived from the intrachromosomal HPV-*TP63* DNAs. These findings define previously unrecognized higher-order architecture of integrated HPV DNA and highlight the power of FISH for distinguishing intrachromosomal from extrachromosomal DNA structures.

**GRAPHICAL ABSTRACT:** 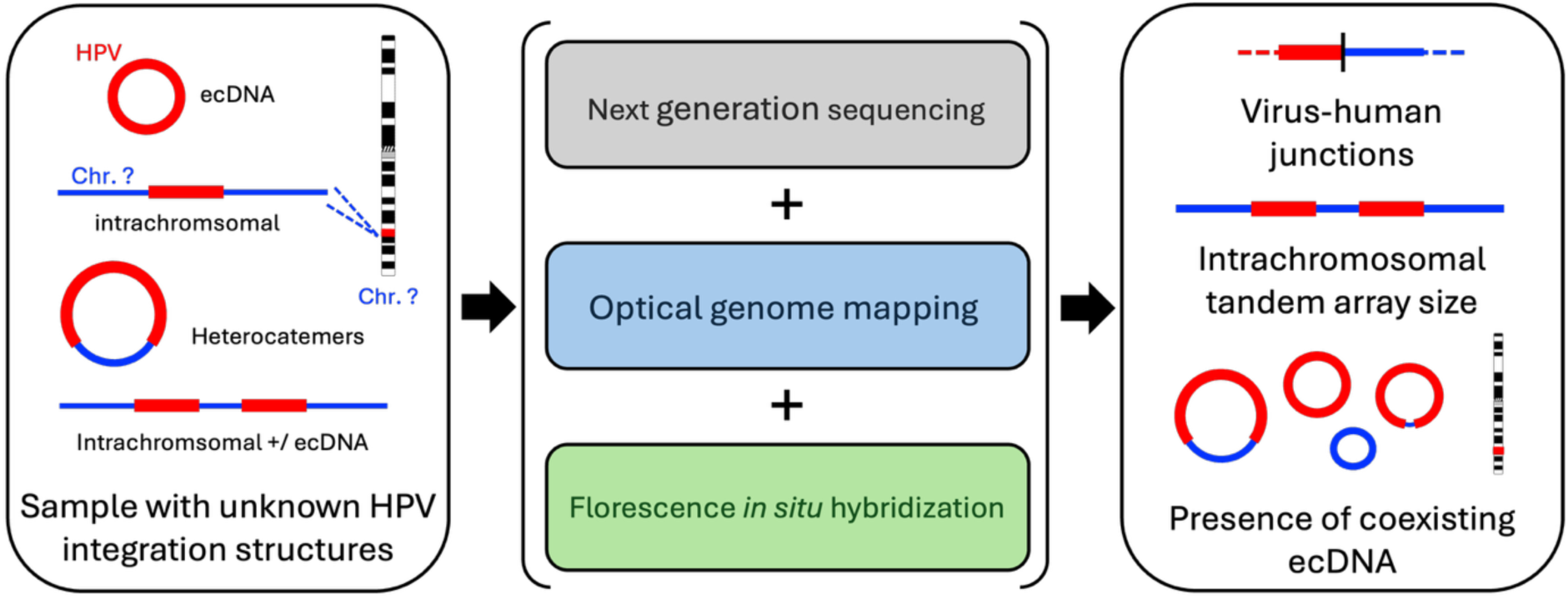

## INTRODUCTION

Human papillomaviruses (HPVs) are a major cause of human cancers, responsible for an estimated 690,000 new cases globally and 38,000 annually in the United States(1,2). Nearly all cervical cancers and approximately 30% of head and neck squamous cell carcinomas (HNSCCs) are associated with an HPV infection(3). During productive infection, HPV genomes replicate as extrachromosomal circular DNA (ecDNA). While most HPV infections are cleared by the immune system, a minority persist and progress to malignancy. In these cases, HPV DNA often becomes integrated with the human genome, typically via aberrant DNA repair, driving clonal expansion of affected cells. Tumor cell clones that emerge consistently retain a segment of the viral genome spanning the upstream regulatory region (URR) and the E6 and E7 oncogenes(4,5). Integration stabilizes these viral oncogenes within the host genome and enables their persistent expression as confirmed by RNA-sequencing(6), leading to inactivation of tumor suppressor genes *TP53* and *pRb*(6–8). DNA sequencing studies revealed structures of the integrated DNAs (4,9–15). These factors are key to HPV-driven oncogenesis(7). HPV DNA integration with human DNA is common in HPV-associated cancers, occurring in at least 83% of cervical cancers (16) and approximately 70% of HPV-positive HNSCC(17–19). Integrated viral DNA is also associated with host genome instability, including structural rearrangements, mutations, and aneuploidy(9,20–22).

It is less certain whether integrated HPV-human DNAs exist as extrachromosomal DNA (ecDNA), intrachromosomal DNA, or both simultaneously. Recent cancer genomics studies have established ecDNA as a key mechanism for oncogene amplification, tumor heterogeneity, and drug resistance across cancer types, including glioblastoma, lung, cervical, and additional cancer types (23–25). ecDNAs can replicate independently, amplify oncogenic content, and contain both viral and host sequences. In HPV-driven tumors, ecDNAs may arise from viral DNA integration events with human DNA, resulting in a hybrid, “heterocatemer” form, but their structural organization and genomic context remain incompletely resolved. Notably, ecDNA circles and chromosomally integrated tandem repeats can share identical sequences, making sequencing-based approaches, whether short- or long-read, insufficient to distinguish them. This distinction is not trivial because intra- and extrachromosomal architectures differ in mechanisms of formation, stability, inheritance, and likely their potential to accelerate tumor evolution. Resolving these fundamental topological ambiguities requires multi-scale approaches that integrate nucleotide-level, structural, and cytogenetic resolution.

Multiple sequencing-based approaches have been used to characterize HPV-human DNA structures in tumors and cell lines, including hybridization capture with short-read sequencing (HC+SEQ), RNA sequencing (RNA-seq), whole-genome sequencing (WGS), long-read sequencing (LRS), and predictive computational tools such as Amplicon Architect(26). HC+SEQ and WGS detect virus-human junctions and have established that integrated HPV DNA typically exists as a sub-genomic segment or as tandem HPV units exceeding one viral genome in length(4,5,9,27), supporting models in which partially replicated HPV DNAs serve as precursors to integrated forms(28). WGS and LRS have revealed tandemly arranged, HPV-human, heterocatemer repeat units(4,9–13). These heterocatemers may be chromosomally associated or exist as extrachromosomal circles, as suggested by methods including E2 to E6 gene ratios, Amplicon Architect analysis of WGS data, LRS, and Fluorescence *in situ* Hybridization (FISH)(4,11,14,15,26,29,30).

These and related approaches have also uncovered widespread genomic instability associated with HPV integration, including structural rearrangements, viral and host mutations, and structural or numerical chromosomal alterations(22,28,31–33). Notably, increased complexity of HPV integration patterns has been correlated with more advanced disease (14,23,34,35) and reduced therapeutic response(36,37). These findings suggest that the architecture of HPV-human DNA integration structures may result from progressive genomic instability during tumor progression and may contribute to oncogenesis.

To elucidate HPV-human DNA structures precisely, we applied a comprehensive battery of genomic and imaging techniques to the HPV16-positive oropharyngeal squamous cell carcinoma cell line UM-SCC-47 (SCC47) as a proof-of-concept system. Using HC+SEQ, RNA-seq, Illumina WGS, Nanopore LRS, optical genome mapping (OGM), and single-cell FISH, we uncovered previously unrecognized structural complexity, including higher-order, multi-megabase architectures of HPV DNA integration, as well as clearly defined chromosomally integrated versus ecDNA structures.

## MATERIAL AND METHODS

### Cell Culture Conditions

The UM-SCC-47 cell line(38) (RRID:CVCL_7759) derived from a squamous cell carcinoma of the oral tongue from a 53Y M was obtained from MilliporeSigma and cultured in Dulbecco’s Modified Eagle Medium (DMEM, high-glucose formulation with sodium pyruvate and L-glutamine; Gibco) supplemented with 10% fetal bovine serum (FBS; Gibco) and 1× non-essential amino acids (NEAA; Gibco). Cells were maintained at 37°C in a humidified incubator with 5% CO₂ and passaged every 3–4 days using 0.5% trypsin-EDTA (Gibco) for detachment.

Cell identity was verified by short tandem repeat (STR) profiling and matched to the reference profile for UM-SCC-47(39). Mycoplasma-free conditions were ensured by routine inspection of DAPI-stained cultures under fluorescence microscopy. Cultures were only used for downstream applications if no extranuclear punctate DAPI signals indicative of mycoplasma contamination were observed.

### Drug Treatment Experiments

For drug treatment experiments, UM-SCC47 and HaCaT cells were seeded at 0.15x10^6^ cells/well in Corning 48-well plates. Cells were placed in the incubator and allowed to adhere overnight. Next, they were treated with various concentrations of (+)-JQ1(Cayman Chemical, Cat. #11187) (0.01μM, 0.03μM, 0.05μM, 0.1μM, 0.3μM, 0.5μM, and 1.0μM). DMSO and untreated cells were used as dual negative controls. Proliferation was monitored using the IncuCyte SX5 live-cell imaging system (Essen BioScience). Phase-contrast images were acquired every 4 hours for 72 hours, and confluence was quantified using IncuCyte software **(version 2024B).** Each condition was run in duplicate.

### Short-Read DNA Hybridization Capture Sequencing

DNA was isolated from approximately 5 x 10^6^ pelleted SCC47 cells using the Quick-DNA Miniprep Kit (Zymo Research, Irvine CA) following the manufacturer’s standard protocol. DNA concentration and purity were assessed using a NanoDrop spectrophotometer.

A custom probe set was designed using the KAPA HyperExplore Custom Capture Panel (Roche Diagnostics, Indianapolis, IN) with tiled coverage across the full-length genome of HPV16(27). Genomic DNA (100-150ng) was mechanically sheared to an average fragment size of 180–220 bp using a Covaris M220 ultrasonicator (Covaris, Woburn, MA). Fragmented DNA underwent end-repair, A-tailing, and ligation of Illumina-compatible adapters using the KAPA HyperPrep Kit (Roche Diagnostics) according to the manufacturer’s protocol. Following pre-capture PCR amplification [8 total cycles], adapter-ligated DNA was hybridized to biotinylated HPV16-specific probes using the KAPA HyperCap workflow (Roche), including post-capture amplification [14 cycles], and bead-based purification.

The captured library was sequenced on a single lane of an Illumina HiSeq system (Illumina, San Diego, CA) to generate paired-end reads. Sequencing reads were processed to remove duplicates, aligned to the human genome (hg38) and HPV reference genomes using the CTAT Virus Integration Finder (CTAT-VIF) pipeline(40). Chimeric reads spanning virus-human junctions were used to identify integration breakpoints. HPV genome annotation and junction visualization were guided by sequence coordinates obtained from the Papillomavirus Episteme (PaVE) database(41).

### Short-Read Whole Genome Sequencing (WGS)

DNA was isolated from approximately 5 x 10^6^ pelleted SCC47 cells using the Quick-DNA Miniprep Kit (Zymo Research, Irvine CA) following the manufacturer’s standard protocol. DNA concentration and purity were assessed using a NanoDrop spectrophotometer.

WGS sequencing and analysis were performed by Novogene under a contract. Following standard quality control procedures, genomic DNA was fragmented to approximately 300bp and used to prepare a single sequencing library. This library was sequenced on the Illumina NovaSeq X Plus platform to generate 2x150 bp paired-end reads. FASTQ files were generated through base calling. The raw reads underwent further quality filtering and were aligned to the human reference genome (hg38) using Burrows-Wheeler Aligner (BWA) software(42). The resulting BAM files were analyzed using the Amplicon Architect pipeline(26) through the AmpliconSuite module on the GenePattern platform(43). Zipped FASTQ files were used as the input, and a custom hg38_viral genome was used for alignment and structural inference.

### Short-Read RNA Sequencing

SCC47 cells were grown to approximately 90% confluence, washed twice with phosphate-buffered saline (PBS), and harvested using TRIzol Reagent (Invitrogen). Lysates were homogenized using QIAshredder (QIAGEN N.V.), and Total RNA was extracted using the RNeasy Mini Kit (QIAGEN N.V.) following the manufacturer’s protocol. Short-read RNA sequencing was performed by Novogene under a contract. Only ribosome-depleted RNA was retained and cleaned by ethanol precipitation prior to library construction.

Sequencing libraries were generated from the rRNA-depleted RNA. After fragmentation, first-strand cDNA was synthesized using random hexamer primers. Second-strand cDNA synthesis was performed using dUTPs in the reaction buffer to enable strand specificity. The directional library was completed through end-repair, A-tailing, adapter ligation, size selection, USER enzyme digestion to degrade the dUTP-containing strand, amplification, and purification.

Final libraries were quantified using both Qubit fluorometry and real-time PCR, and fragment size distribution was assessed using a Bioanalyzer. Quantified libraries were then sequenced on the Illumina NovaSeq 6000 platform. Resulting BAM files were uploaded to the Integrative Genomics Viewer (IGV) for visualization and analysis.

### Long-Read DNA Sequencing

High molecular weight genomic DNA was isolated from approximately 8x10^6^ pelleted SCC47 cells using the Gentra Puregene Cell Kit (QIAGEN N.V.), with protocol modifications optimized for LRS. Centrifugation steps were performed at increased speeds up to 24,000 x g whenever possible to improve pellet quality. Cell lysis was carried on with RNAseA at 37°C, and the lysate was incubated for 50 minutes to ensure complete DNA degradation. Protein precipitation was conducted on ice for 10 minutes with minimal vortexing to preserve DNA integrity. DNA was then precipitated, and the resulting pellet was washed thoroughly with 70% ethanol before air-drying. DNA was eluted in 200 µL volume of 1M Tris-HCl (ph 7.5). DNA yield was quantified using the Qubit 3.0 Fluorometer (Thermofisher, Waltham MA) and the Qubit dsDNA High Sensitivity kit (Thermofisher, Waltham MA). DNA fragment lengths were assessed using the Femto Pulse system (Agilent, Santa Clara CA) and the gDNA 165 kb kit. Contaminant carryover and RNA co-purification was assessed via Nanodrop spectrophotometry by examining 260:230 and 260:280 ratios. High protein carryover in the

DNA isolate was enzymatically removed using thermolabile proteinase K (NEB #P8111S, Ipswich MA) in a 90 µL reaction with Q5 reaction buffer (NEB #B9027S, Ipswich MA). DNA quality after cleanup was reevaluated using the Femto Pulse, Qubit Fluorometer, and Nanodrop platforms.

After QC 3.4 µg of purified genomic DNA was used to prepare a library for sequencing using the ligation kit (SQK-LSK110, ONT), following the manufacturer’s instructions with specific modifications. The FFPE and DNA repair step was extended to 1 hour, and adapter ligation was performed for 1 hour. The final library was eluted in 34 µL of elution buffer.

For sequencing, 11 µL of the DNA eluate library containing 25.4 fmol of input (assuming a mean fragment length of 50 kb) was loaded onto a FLO-PRO-0002 PromethION flowcell (R9.4.1). Initial sequencing was conducted for 22 hours. The flow cell was then washed using the Flow Cell Wash Kit (EXP-WSH004, ONT) following the manufacturer’s instructions and reloaded with an additional 25.4 fmol of the same library. A second round of sequencing ran for 21 hours, after which the flow cell was washed and reloaded a final time with 25.4 fmol of library. Sequencing preceded for an additional 43 hours, resulting in a total of 167.24 GB and 152 million reads.

Raw sequencing data was basecalled using guppy version 6.3.2 (ONT). Basecalled reads were aligned to the human reference genome (hg38) using Minimap version 2.24, optimized for long-read mapping.

To annotate viral and chimeric integration events, a custom nucleotide sequence database was assembled containing annotated HPV16 open reading frames and human *TP63* exon sequences. Long reads were queried against this reference using BLASTN(44,45). Alignment features and strand orientation were used to classify and annotate HPV-*TP63* chimeric structures, including junction breakpoints, repeat boundaries, and gene fusions.

ONT genome sequencing reads aligning proximal to the reference genome and proximal to gene *TP63* (overlapping region chr3:189,700,000-190,100,000, hg38) were extracted, totaling 4736 reads. These reads were assembled using the Flye assembler(46) as follows: “flye – nano-raw reads.fq –out-dir flye.outdir –keep-haplotypes”.

### Long-Read RNA Sequencing

RNA was prepared from SCC47 cells as described above. RNA quality was assessed using the Agilent 2100 Bioanalyzer with the RNA 6000 Nano kit, and concentration was quantified using the Qubit (Thermofisher, Waltham MA). Approximately 344 ng of high-quality RNA was used for cDNA synthesis and library preparation using the SQK-PCS110 kit (ONT) following the manufacturer’s instructions. Based on assumed mean fragment length of 1.5 kb, 230 fmol of adapter-ligated cDNA was eluted into 24 µL of elution buffer (ONT). For sequencing, 12 µL of the cDNA eluate library was loaded one FLO-PRO-002 PromethION flowcell (R9.4.1) and sequenced continuously for 89.3 hours, generating 119.9 GB of data and approximately 9.87 million reads.

ONT transcriptome sequencing matching the HPV16 genome were identified via minimap2 alignment(47), yielding 14,005 transcriptome reads. These reads were *de novo* assembled using RATTLE(48). Commands were as follows:

rattle cluster -I hpv16_matching_reads.fq -t 1 -o .
rattle cluster -I hpv16_matching_reads.fq –iso -t 1 -o .
rattle cluster_summary -I hpv16_matching_reads.fq -c clusters.out
mkdir clusters
rattle extract_clusters -I hpv16_matching_reads.fq -c clusters.out -o clusters –fastq
rattle correct -I hpv16_matching_reads.fq -c clusters.out -t 4 2>&1 | tee
rattle.correct.log
rattle polish -I consensi.fq -t 8 –rna

### Optical genome mapping

Ultra-high molecular weight DNA was extracted from 1.5 million SCC47 cells using the Bionano Prep SP-G2 Blood & Cell Culture DNA Isolation Kit (PN 80060). Briefly, cells that had been previously cryopreserved as a dry pellet and stored at -80°C were pelleted at 500xg for 5 minutes at 4°C, then washed in 1mL DNA stabilizing buffer. Cell counts were verified, and 1.5 million cells were centrifuged again for 2 minutes at 2200 xg at 4°C to create a compact pellet. DNA was isolated from this compact cell pellet following Bionano Prep SP-G2 Frozen Cell Pellet DNA Isolation Protocol, rev C. This DNA was fluorescently labeled at the recognition site CTTAAG with the enzyme DLE-1 and counter-stained, using the Bionano Prep DLS-G2 Labeling Kit (PN 80046) following Bionano Prep DLS-G2 Protocol, rev E. Fluorescent imaging of single molecules was performed on the 2nd generation Bionano Saphyr system with instrument control software (ICS) version 5.3.23013.1, collecting approximately 300X effective coverage of the reference hg38 after automatic filtering for noise and molecule quality. OGM data was assembled using the Guided Assembly algorithm with default parameters for low allele fraction SV-calling using Bionano Access version 1.8.2.

### Fluorescence *In Situ* Hybridization (FISH)

SCC47 cells were grown to 60-80% confluence and treated with 0.1 □g/ml colcemid (ThermoFisher, Waltham, MA) to arrest cells in metaphase. Floating (shake-off) cells were collected, and adherent cells were detached by trypsinization. JQ1-treated cells were harvested after 48 hours of exposure to 1μM, 0.05μM, or 0.01μM of JQ1 or vehicle control. Adherent cells in metaphase were obtained by KaryoMAX (Gobco) treatment at 0.1mg/mL for 30 min. Cells were washed with PBS and trypsinized, and cell were collected.

All cells were exposed to pre-warmed (37°C) 0.075M potassium chloride as a hypotonic treatment. Following incubation, cells were pelleted, fixed in methanol and acetic acid (3:1), and washed twice with the same fixative. Cell suspensions were dropped onto slides to generate both metaphase spreads and interphase nuclei. Separately, cells were grown directly on a glass coverslip (22x22 mm^2^) and fixed using the same 3:1 methanol:acetic acid solution to allow *in situ* hybridization of interphase nuclei.

Fluorescence *In Situ* Hybridization (FISH) was performed to visualize the HPV16 integration using protocols described previously(49). Slides were baked at 95°C for 5 min and prewashed in 2xSSC at 72°C for 2 min. Slides were then incubated for 8-min with 5 μl of pepsin solution (100 mg/ml in 0.01 N HCl) to facilitate probe access to the nucleus. Custom DNA probes were prepared to target specific genomic regions of interest: *TP63*: BAC RP11-373I6 (chr3:189,779,579–189,959,633; includes 3′ end with tandem repeat in SCC47); TERC: BAC RP11-990E14 (chr3:169,692,858–169,870,368); HPV16: Full-length HPV16 plasmid (source: Van Doorslaer et al.(50,51)). DNA from BACs and HPV16 plasmid was extracted and labeled by nick translation as previously described(52,53), using Dyomics modified dUTPs: DY-495-aadUTP (Spectrum Green) for RP11-990E14 (TERC), DY-530-aadUTP (Spectrum Gold) for RP11-373I6 (*TP63*), and DY-590-dUTP (Spectrum Red) for HPV16. Probes and slides were co-denatured at 75°C for 3 min and hybridized overnight at 37°C using the ThermoBrite System (Abbott Molecular, Des Plaines, IL). Post-hybridization washes included 0.4X SSC/0.3% NP-40 at 74°C for 2-4 min, followed by 2X SCC/0.1% NP-40 at room temperature for 2 min. Slides were dehydrated via ethanol series and mounted with 30 μl of VECTASHIELD antifade mounting media containing DAPI (Vector laboratories INC.), covered with a coverslip.

The vulva tumor sample was obtained from The Cooperative Human Tissue Network (CHTN) and processed for FISH analysis as previously reported(4). Briefly, tissue sections of ∼5–7 μm in thickness were preserved in Optimal Cutting Temperature compound and were fixed in ice-cold methanol for 10min. A 10-min incubation with 30 μl of pepsin solution (100 mg/ml in 0.01N HCl) was used to facilitate the subsequent hybridization of custom DNA probes. Custom DNA probes were prepared as above to target specific genomic regions of interest: RP11-662P23 (hg38 -chr5:56,420,151-56,608,998 mapping to the specific region of HPV16 integration on chromosome 5 in Tumor 4); Full-length HPV16 plasmid (source: Van Doorslaer et al.(50,51)). BAC DNA and HPV16 plasmid DNA were isolated and labeled by nick translation as above using the following dUTPs from Dyomics (Jena, GE, USA): DY-495-dUTP (within the green spectrum to visualize RP11-662P23) and DY-590-dUTP (within the red spectrum to visualize HPV16). Probes and slides were denatured simultaneously at 74 °C for 5 min and hybridized at 37 °C overnight using the ThermoBrite System (Abbott Molecular). Post-hybridization washes included 0.4X SSC/0.3% NP-40 at 74°C for 2 min, followed by 4X SCC/0.1% Tween-20 at room temperature for 3 min. Slides were dehydrated via ethanol serial washing and mounted with 50 μl of VECTASHIELD antifade mounting media containing DAPI (Vector laboratories INC.), covered with a coverslip. The experimental procedures were approved by the Internal Review Board of the Rutgers Cancer Institute (IRB#:002215). The human studies conducted in this research were performed in full compliance with all relevant ethical regulations, including adherence to the principles outlined in the Declaration of Helsinki.

FISH imaging was performed using the BioView Duet scanning system (BioView USA Inc., Billerica, MA) mounted on an Olympus BX63 microscope (Olympus, Tokyo, Japan) equipped with a 60x oil-immersion objective (NA 1.35). For each image, 11 Z-stack focal planes (0.5 μm per step) were captured to ensure detection of signals across depths. Fluorescent emissions were recorded using the following Semrock Optical Filters (IDEX Health & Science, LLC; Rochester, NY): DAPI: 415–475 nm; Spectrum Green (TERC): 519–537 nm; Spectrum Gold (*TP63*): 558–587 nm; Spectrum Red (HPV16): 609–643 nm. Images of metaphase and interphase nuclei were acquired in both combined and individual color channels. Signal counts from BioView AI-assisted scoring were manually curated and quantified by hand in and around each nucleus.

### Statistical Analysis

Fisher’s exact test was used to evaluate the difference in frequency of nuclei containing both intrachromosomal and ecDNA between interphase nuclei prepared under isotonic versus hypotonic conditions. Student’s T-test was applied to compare the *TP63* signal size between copies of chromosome 3 with HPV16 intrachromosomal integrations and those without HPV16 signals, and to compare total normalized object count between control and JQ1-treated cells after 48 and 68 hours. The non-parametric Mann-Whitney U test was used to compare the HPV16 signal size between control and JQ1-treated cells due to data not meeting the normality requirement as assessed by Kolmogorov-Smirnov and Shapiro-Wilk tests. The non-parametric Wilcoxon signed ranks test was used to assess differences in signal number between paired loci within each hypotonic interphase nucleus and integrated HPV16 copies between control and JQ1-treated cells due to data not meeting the normality requirement as assessed by Kolmogorov-Smirnov and Shapiro-Wilk tests. All statistical tests were two-sided, and significance was defined as p < 0.05, where a test value less than 0.05 indicated a significant difference. In all cases N represents the number of cell nuclei analyzed. Cell nuclei that could not be clearly visualized due to significant overlap with another nucleus or not being completely included in the field of view were excluded. Analyses were performed using GraphPad Prism 10.4.2.

## RESULTS

### Structure of the integrated HPV16 DNA segment in SCC47 determined by hybridization capture and short-read sequencing (HC+SEQ)

To map the viral-human DNA junctions in the SCC47 cell line, we performed hybridization capture followed by short-read sequencing. HPV-DNA-containing fragments were enriched from sheared total genomic DNA using a custom probe set targeting the complete HPV16 genome as previously described(4). Illumina sequencing, coupled with computational analysis using CTAT-VIF(40), enabled precise mapping of HPV-human junctions and definition of the integrated subgenomic segment at single-nucleotide resolution (**Fig. 1A**).

**Figure 1.**
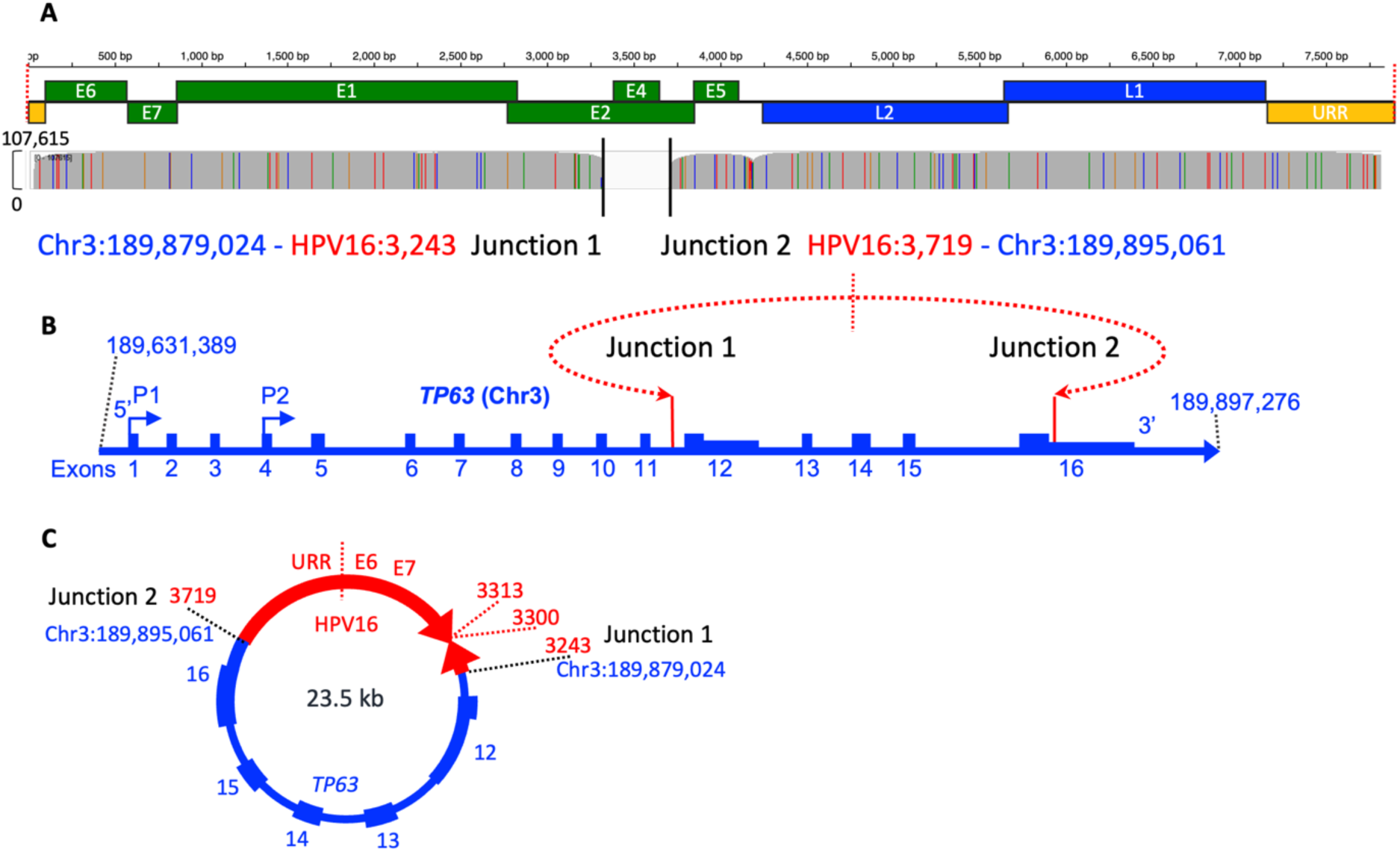
Structure of the HPV16 genome and virus-human DNA junctions revealed by hybridization capture and Illumina sequencing: **A)** Hybridization capture plus sequencing coverage across the HPV16 genome. HPV16 reference genome coordinates and arbitrarily linearized open reading frame map are shown at the top. Deduplicated Illumina read counts are plotted below, with color-coded vertical lines marking single-nucleotide polymorphisms (A=green, T=red, C=blue, G=orange). Black lines labelled Junction 1 and Junction 2 mark the positions and coordinates of the two HPV16-human DNA junctions. B) Diagram of the initial HPV16 DNA integration into the *TP63* gene on human chromosome 3. Transcription start sites at *TP63* exons 1 and 4 are indicated. The HPV16 segment ends are oriented towards each other (red arrows), consistent with a non-linear (circular) integration event. Short vertical lines indicate the junctions between HPV and human DNA. **C)** Schematic of the extrachromosomal circle formed by the integration event shown in panel B. This heterocatemer consists of HPV16 DNA (red) joined to *TP63* exons 12 to 16 (blue). The HPV16 portion lacks a 406 bp segment (positions 3,243-3,719) and includes a 57 bp inverted duplication between positions 3,243 and 3,300.

The integrated HPV16 DNA segment contained nearly the entire viral genome, except for a 406 base pair deletion within the *E2* gene, spanning positions 3,243 to 3,719 (Papillomavirus Episteme, PaVE(41)). Despite an average coverage of 78,550 reads across the HPV16 genome, no reads mapped within this gap, implying that all HPV16 copies in SCC47 lack this segment. The viral DNA was joined to the *TP63* gene on chromosome 3 at hg38 positions 189,879,024 (Junction 1) and 189,895,061 (Junction 2) (**Fig. 1A**). The orientations of the HPV16 and human DNA segments across each junction revealed that the viral ends are directed toward each other, indicating that the original integration event likely formed a circular, extrachromosomal intermediate (**Fig. 1B-C**). Such structures have been referred to as non-linear integrations, in contrast to events where viral ends are oriented away from each other and linearly insert into a human chromosome(54). Junction 1 mapped to intron 11, just upstream from exon 12 of *TP63*, while Junction 2 was within the 3’ untranslated region of exon 16, encompassing 16,037 bp (16 kb) of the 3’ portion of the gene (**Fig. 1B**). Additionally, a 57 bp inverted duplication of HPV16, sequence (positions 3,244 to 3,300) was present at one end of the integrated segment (**Fig. 1C**) for a total of 7557 bp (7.5 kb). The combined HPV16 and human sequences formed a 23,594 (23.5 kb) structure, hereafter termed a heterocatemer, corresponding to the amplified genomic segment previously reported by Akagi et al.(9).

### Tandem repeats of the 23.5 kb HPV16-human DNA heterocatemer identified by LRS

Long-read DNA sequencing of SCC47 cells generated 152 million sequence reads, with 4736 reads mapping to a 400 kb region spanning the *TP63* locus. Assembly with Flye (46) confirmed the exact 23.5 kb HPV16-*TP63* heterocatemer structure predicted by hybridization capture, including both virus-human junctions and the 57 bp inverted duplication (**Figs. 2** and **1B**). All copies lacked the same 406 bp stretch in the E2 gene identified by HC+SEQ, suggesting that all copies of HPV16 in this cell line derived from the original HPV-human DNA integration event described in **Fig. 1**.

**Figure 2.**
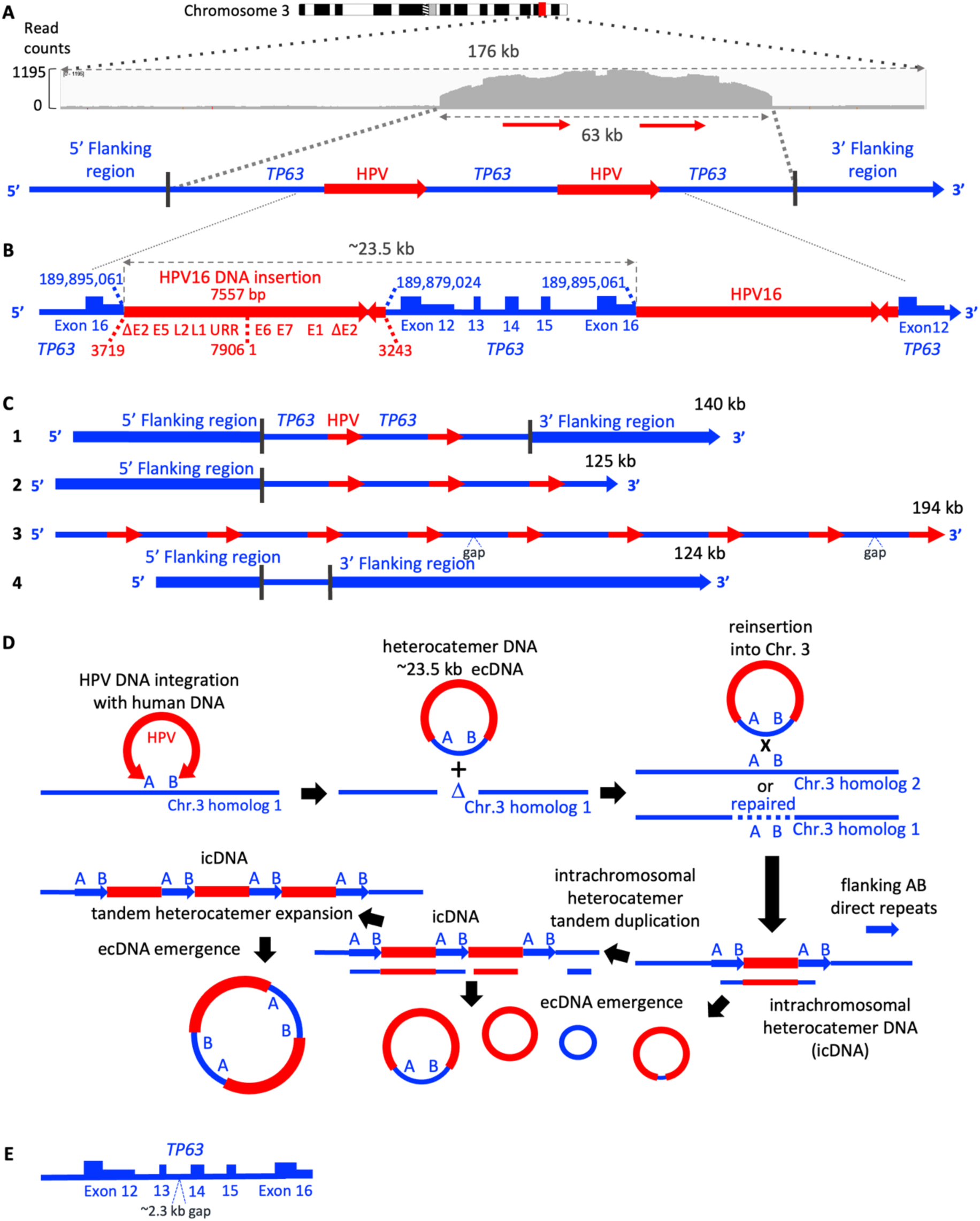
Tandem repeat structures by LRS and a model for their formation: **A)** LRS coverage across a 176 kb region encompassing a 63 kb segment with two tandem units of HPV16 DNA and *TP63* exon 12-16. **B)** Schematic of a single 23.5 kb tandem heterocatemer unit composed of HPV16 DNA (red) and *TP63* exons 12-16 DNA (blue). **C)** Representative long reads showing the HPV-*TP63* heterocatemer structure. Three reads contain variable numbers of tandem repeats (1 to 3); one read contains only normal *TP63* sequence (4). **D)** Proposed model for heterocatemer formation and expansion. A and B identify the *TP63* exon 12-16 segment. An HPV16 DNA segment, potentially from a bidirectional replication intermediate, integrates with its ends oriented toward each other. This configuration leads to: (1) formation of an extrachromosomal circular DNA containing the HPV16 segment and *TP63* exons 12–16 (the AB segment), and (2) a corresponding deletion of the AB segment from one human chromosome. Homologous recombination between the circular intermediate and a second human chromosome (or a repaired version of the original) results in intrachromosomal insertion of the HPV DNA flanked by AB direct repeats. These repeats facilitate amplification through homology-dependent mechanisms, including mitotic DNA polymerase slippage or non-allelic homologous recombination. Additional extrachromosomal circles can form through recombination between ends of incompletely replicated tandem units. **E)** Map of a recurrent ∼2.3 kb deletion within the *TP63* segment observed in some heterocatemer units (e.g., map 3 in panel C), likely arising from a single deletion event propagated by repeat amplification

To quantify the tandem repeats, we constructed a chimeric reference sequence containing two copies of the 7.5 kb HPV16 segment with three 16 kb *TP63* segments, plus flanking *TP63* sequences (**Fig. 2A**). A more detailed structure of the tandem heterocatemers is shown in **Fig. 2B**. Nanopore reads aligned to this reference, visualized in Integrated Genome Viewer (IGV) (55), revealed amplification of heterocatemers relative to background genomic DNA (**Fig. 2A**). Representative genetic maps (**Fig. 2C**) showed variable numbers of tandem 23.5 kb repeats. LRS reads also detected the normal *TP63* allele, which presumably descended from the chromosome 3 homolog where HPV16 DNA did not integrate. A 140 kb read matching the hg38 reference sequence further validated the inferred structure.

Among the 20 longest reads, the maximum number of tandem heterocatemer units was nine, with a modal repeat number of five (**Supplementary Fig. 1**). Exact repeat numbers could only be determined for reads containing *TP63* flanking sequences on both sides of the heterocatemers. Many of the longest reads lacked both flanking sequence (**Supplementary Fig. 2**), meaning their repeat counts are minimum estimates. In reads with flanking human DNA on both sides, the maximum repeat count was two (**Fig. 2C** and **Supplementary Fig. 2**), suggesting that these were among the smaller tandem arrays in SCC47 cells. While LRS could not distinguish whether these structures were from linear intrachromosomal insertions or large ecDNA circles, it revealed extensive and variable heterocatemer repeat instability.

A potential mechanism for the formation of these structures is shown in **Fig. 2D**. Initial integration likely produced a circular heterocatemer composed of HPV and *TP63* sequences. Homologous recombination with chromosomal *TP63* could have inserted this unit intrachromosomally, producing an HPV DNA insertion flanked by direct repeats of the *TP63* segment. Further homology-driven rearrangements involving the flanking direct repeats could then generate arrays of tandem heterocatemers, either as circular or linear forms (**Fig. 2D**).

Additionally, a subset of the 23.5 kb heterocatemer units contained a 2.3 kb identical internal deletion located between *TP63* exons 13 and 14 (**Fig. 2E** and **Supplementary Fig. 2**). This deletion likely occurred once and was propagated in a subset of units by the same large-scale recombination events responsible for tandem repeat expansion and contraction. In summary, LRS revealed amplification of 23.5 kb HPV16-*TP63* heterocatemers in tandem arrays and suggested an ongoing process of recombination-driven structural evolution in SCC47.

### Splice junctions and polyadenylation sites identified by short-and long-read RNA-seq are consistent with transcription from tandem repeat or circular heterocatemer templates

To assess how the SCC47 HPV-human DNA structures identified by LRS influence viral gene expression, we performed both short- and long-read RNA sequencing. Short-read RNA-seq coverage across the HPV16 genome (**Fig. 3A)** confirmed the absence of reads mapping to the 406 bp E2 deletion, consistent with the integrated DNA structure. Transcripts containing the E6 and E7 open reading frames (ORFs) were detected, initiating near the viral early promoter upstream of the E6 ORF. The E6*I splice variant, an out-of-frame event generated by splicing from positions 226 to 409 and common in HPV-positive tumors(4,56,57), was also present (**Fig. 3A**). Transcripts from the E1 ORF and the truncated, inversion-containing E2 gene were detected at lower levels, possibly because they reside within intronic regions of sequences that continue further downstream.

**Figure 3.**
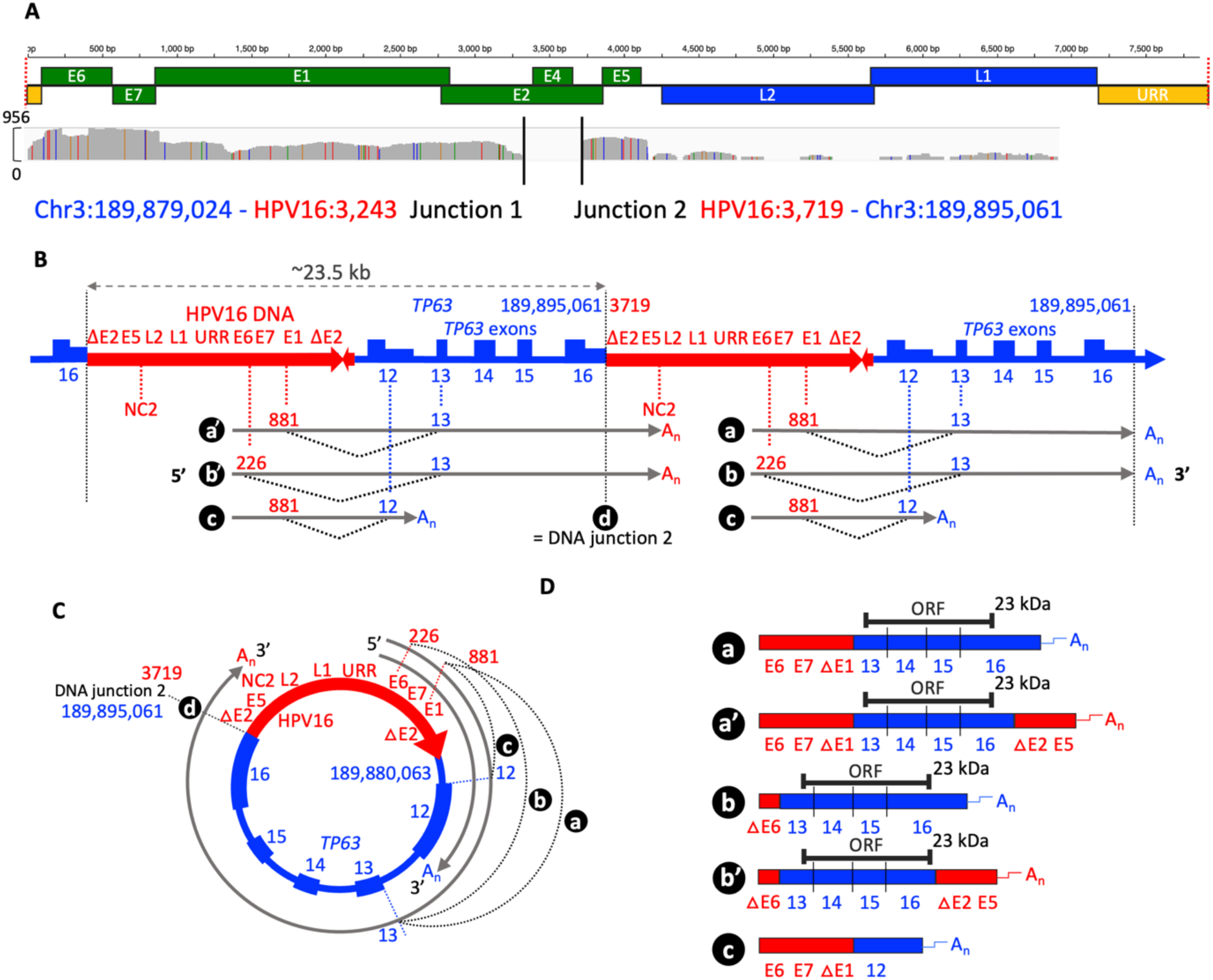
Characterization of HPV16-*TP63* fusion transcripts, splicing patterns, and polyadenylation sites: **A)** RNA-seq read coverage across the HPV16 genome reveals transcriptional activity and confirms the 406 bp E2 deletion (absence of reads across that region). Solid black lines indicate the two HPV16-human DNA junctions. **B)** Splicing patterns and predicted transcripts identified by long-read RNA-seq are consistent with short-read RNA-seq. HPV16 genomic sequences (red) and *TP63* exons 12–16 (blue) are shown with primary transcripts in gray. Specific variably spliced and variably polyadenylated primary transcripts are identified by a, a’, b, b’, and c with splice junctions indicated by dotted gray lines. Splicing junctions (dotted gray lines) connect viral 5’ splice donor sites to human 3’ splice acceptors, generating five distinct fusion transcript isoforms: a, a′, b, b′, and c. Polyadenylation sites are mapped to either HPV16 NC2 (red) or *TP63* exons 12 or 16 (blue). Site d denotes transcripts spanning Junction 2 in transcripts a′ and b′. **C)** Schematic showing how transcripts a, b, and c could arise from transcription of an extrachromosomal circular heterocatemer DNA template. **D)** Spliced RNA isoforms with HPV16 sequences in red and *TP63* sequences in blue. A predicted open reading frame (black) encodes a ∼23.5kDa *TP63-*derived protein from transcripts b and b′, initiating at an internal AUG in exon 13 and terminating in exon 16.

Chimeric viral-host transcripts were detected with splice patterns that implied they originated from heterocatemer templates. Two major 5’ splice donor sites (5’ss) in HPV16 were joined to 3’ splice acceptor sites (3’ss) in *TP6*3 (**Fig. 3B**). The most prominent 5’ss was the E1^E4 5’ss at position 881 near the E1 ORF start that is typically used in HPV-transformed cells(4) and which spliced to 3’ss of *TP63* exons 12 or 13. Interestingly, the 406 bp E2 deletion removed the E1^E4 3’ss, presumably allowing splicing to the *TP63* exons. Another 5’ss at position 226 within E6 spliced to the 3’ss of *TP63* exon 13. These spliced products are consistent with transcription from either tandem heterocatemer arrays or circular ecDNA heterocatemers (**Fig. 3B-C**). Curiously, RNA reads also contained the E5 ORF (**Fig. 3A**), which is located upstream of the URR transcription initiation site in the integrated 7.5 kb HPV DNA segments (**Fig. 3B**).

Additionally, RNA-seq reads were detected bridging *TP63* exon 16 into HPV16 E2, aligning exactly with DNA Junction 2, the site of HPV16 DNA integration. This indicated that transcription proceeds across the heterocatemer from *TP63* into HPV16, which is likely because exon 16 in the heterocatemers is truncated before its polyadenylation site, thereby allowing transcription to continue downstream into viral sequences including the E5 ORF (**Figs. 3B-C**).

To validate these structures and determine polyadenylation site usage, we performed long-read Nanopore RNA-seq. This confirmed the splicing patterns described above and identified three utilized polyadenylation sites: the HPV non-coding region 2 (NC2) site downstream of E5, *TP63* exon 12, and *TP63* exon 16 (**Fig. 3D**). During productive HPV infection, E6/E7 mRNAs are typically polyadenylated at NC2(58). In cancer, however, polyadenylation often occurs at host sites following viral-host splicing events(59,60), consistent with our findings (**Figs. 3B-C**).

*TP63* itself is known to undergo alternative splicing and polyadenylation at sites in either exon 12 or exon 16, depending on alternative isoform expression(61,62). Transcripts from SCC47 heterocatemers (**Fig. 3B**), whether chromosomally integrated or ecDNA circles, initiate within an HPV segment and proceed into the downstream *TP63* sequence. Those spliced to exon 12 are polyadenylated at that site. Those alternatively spliced to exon 13 proceed through the *TP63* segment and into the next HPV segment, where they are polyadenylated at NC2. However, transcripts initiating from the HPV segment in the final unit of a chromosomally integrated tandem array are polyadenylated at the intact exon 16 site (**Fig. 3B**).

The mature, spliced, and polyadenylated transcripts identified by long-read RNA sequencing are shown in **Fig. 3D**. A small *TP63* protein of approximately 26 kDa was previously observed in SCC47 cells using anti-*TP63* antiserum in western blot analyses(9). A candidate ORF consistent with this size begins at an internal AUG codon in exon 13 and remains in-frame through the canonical stop codon in exon 16. This ORF would encode a predicted 23.5kDa protein composed entirely of *TP63-*derived sequence. It could be translated from two spliced, monocistronic transcripts we identified (mRNAs b and b’, **Fig. 3D**).

In conclusion, the RNA structures identified in SCC47 include specific HPV-*TP63* chimeric transcripts generated by alternative splicing and variable polyadenylation site usage. These transcripts are consistent with the underlying HPV16-human DNA structures and could encode viral E6 and E7 oncoproteins. Transcripts polyadenylated at the exon 16 site could originate only from templates that included *TP63* DNA flanking heterocatemer array, arguing that at least some of the transcripts originated from intrachromosomal DNA. Although speculative, two transcripts may also encode a truncated *TP63* protein composed solely of the carboxyl-terminal exons of the gene.

### *TP63* rearrangements determined by WGS and Amplicon Architect

The LRS and RNA-seq analyses identified heterocatemer tandem repeat and ecDNA structures and how they allow transcription of the HPV16 oncogenes, but determination of the large-scale architecture of the HPV-human DNA structures required additional genomic tools. First, WGS and computational analysis using Amplicon Architect(26) were used to characterize structural rearrangements at the *TP63* locus. WGS reads visualized in IGV (**Fig. 4A**) confirmed the features identified by HC+SEQ and LRS, including HPV16 coverage across the integrated region, the 406 bp deletion, and both HPV-human junctions (**Fig. 1A** and **Fig. 2**).

**Figure 4.**
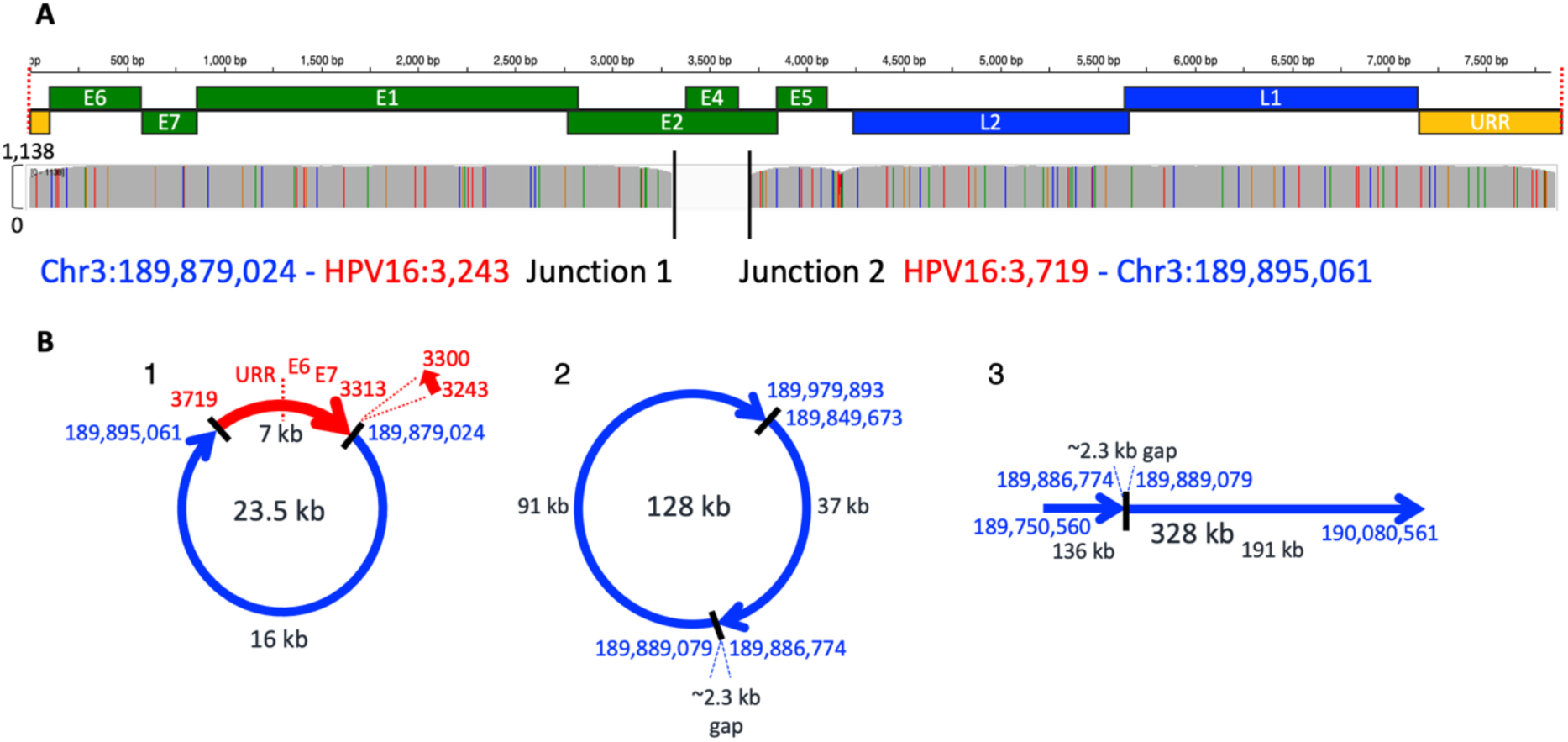
Identification of DNA junctions by WGS and Amplicon Architect: **A)** WGS read coverage across the HPV16 genome. Black lines indicate the two HPV16-human DNA junctions. **B)** Predicted DNA structures from Amplicon Architect analysis. HPV16 segments are shown in red and human *TP63* sequences in blue; junctions are indicated by black bars. Structure 1 recapitulates the canonical 23.5 kb HPV-*TP63* heterocatemer defined by HC+SEQ and LRS, including the characteristic E2 deletion and 57 bp inversion. Structure 2 is an alternative ecDNA-like circle derived from junctions within *TP63* and may represent circularization of rearranged segments. Structure 3 depicts a linear DNA configuration containing the ∼2.3 kb *TP63* deletion found in a subset of tandem repeat units (also identified by LRS).

Amplicon Architect aligned reads to a hybrid reference genome (hg38 and HPV16) and performed copy-number-aware breakpoint analysis to identify connected amplified segments. Prediction 1 (**Fig. 4B**) recapitulated the exact 23.5 kb HPV16-*TP63* heterocatemer defined by HC+SEQ and long-read genomic sequencing, including the absence of the 406 bp E2 deletion and the 57 bp inverted duplication. Although this structure was classified as ecDNA-like, it could also represent a tandem linear unit with identical sequence architecture, highlighting a limitation of short-read data in resolving DNA topology.

Prediction 2 revealed an alternative ecDNA-like configuration involving two amplified *TP63*-derived junctions formed by breakage and rejoining of segments within *TP63* (**Fig. 4B**). One junction matched the precise 2.3 kb deletion observed by long-read DNA sequencing (**Fig. 2E** and **Supplementary Fig. 2**) and was included in Prediction 3. The second junction connected flanking regions of the HPV16 integration site within *TP63* at chr3:189,849,673 and chr3:189,979,893 (**Fig. 4B**), likely reflecting a large-scale rearrangement involving sequences flanking the tandem repeat arrays.

In summary, WGS and Amplicon Architect confirmed the major features of the HPV16-*TP63* integration, including structural variants and breakpoints, but did not resolve the higher-order tandem repeats. The results also revealed additional large-size-scale human genomic rearrangements flanking the integrated heterocatemers.

### Megabase-scale organization of HPV-driven rearrangements defined by OGM

Determining the exact number of tandem units and the full extent of higher-order organization of these structures exceeds the resolution limits of even the longest-read sequencing technologies. To resolve these larger-scale features, we employed OGM, which enables visualization of genomic architecture at the megabase scale. OGM uses fluorescent labeling at the specific DNA motif CTTAG, recognized by the Direct Label Enzyme (DLE-1), to image long, linearized individual DNA molecules. The resulting labeled molecules are computationally aligned to the human genome based on their unique DLE-1 site patterns and assembled into consensus sequence maps. Because the integrated HPV16 segment in SCC47 lacks DLE-1 recognition sites, it appears as a gap in the labeling pattern that spans the 7.5 kb length viral segment plus the length of the adjacent human sequences extending to the nearest flanking DLE-1 sites.

An OGM assembly aligning to the *TP63* locus identified two insertions marked by red-labeled DLE-1 sites (**Fig. 5A**). Closer inspection of one insertion revealed a repeating pattern of ∼9 kb gaps (labeled 1 through 10), corresponding to ten HPV DNA units (∼ 7.5 kb each) flanked by human sequence extending to the nearest DLE-1 sites (**Fig. 5A**). These units were interspersed with ∼14 kb segments of *TP63*, consistent with the 23.5 kb total heterocatemer unit size inferred from LRS. Thus, OGM determined the number of heterocatemers per array and defined the human sequences flanking each tandem repeat array.

**Figure 5.**
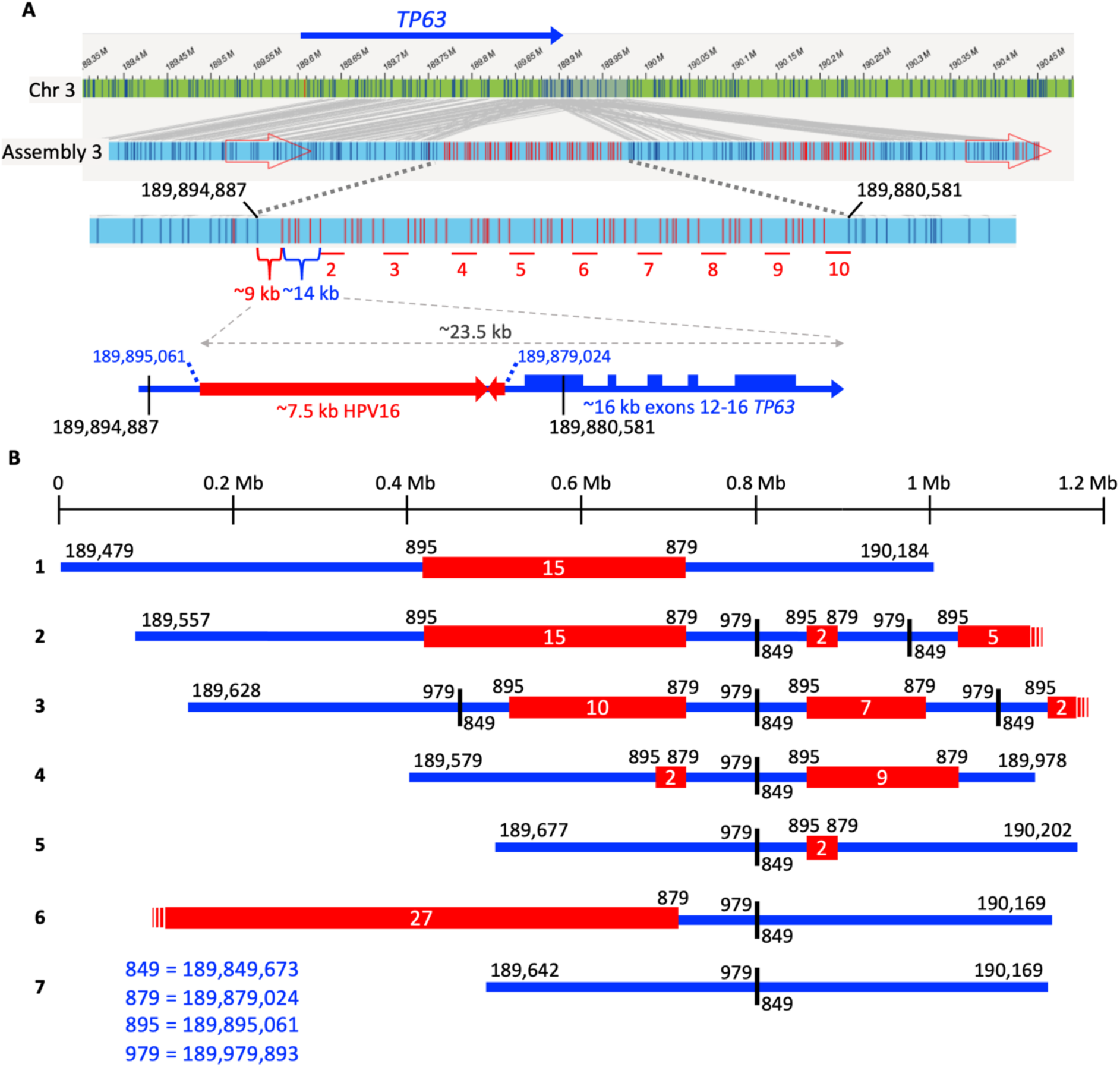
Megabase-scale structures of HPV16-*TP63* tandem repeats revealed by optical genome mapping: **A)** OGM mapping of SCC47 cells identified a large *HPV-TP63* heterocatemer tandem repeat insertion (Assembly 3) within chromosome 3. The *TP63* region of chromosome 3 is shown in green with DLE-1 labeling sites (blue vertical lines) and hg38 genomic coordinates. Assembly 3 (light blue) is aligned to chromosome 3 (gray connecting lines), with two large insertions (gaps in labeling pattern) corresponding to HPV-human heterocatemer tandem repeats. Vertical red lines mark DLE-1 sites located within *TP63* exons 12-16. The blue rectangle below highlights one insertion in detail, showing a repeating 23.5 kb unit composed of ∼14 kb *TP63* sequence (with red DLE-1 sites) and ∼9 kb HPV16 DNA lacking DLE-1 sites, bounded by ∼2 kb of *TP63* sequence to the nearest DLE-1 sites. Ten such tandem heterocatemers are present in this array. A genetic map of a single 23.5 kb unit is shown for reference. **B)** Summary of all seven OGM assemblies (1–7). Megabase size scale is shown above. Each assembly includes sections aligning fully to the human genome (blue) and interspersed tandem arrays of HPV16-*TP63* heterocatemers (red). Tandem unit copy numbers are shown in white text. Human-human rearrangement junctions are marked with vertical black bars. DLE-1 site coordinates at the start and end of each assembly are shown using a simplified six-digit notation, omitting the final three digits. Assemblies beginning or ending within tandem arrays are indicated by thin red vertical lines. The middle three digits of the human genome coordinates are used to label array and junction boundaries. A key listing the full human genomic positions is shown in the bottom left corner.

OGM identified seven distinct assemblies, ranging from ∼0.6 Mb to ∼1.2 Mb in size (**Fig. 5B** and **Supplementary Fig. 3**), each with a unique pattern of heterocatemer tandem repeats ranging from a single unit to at least 27 in Assembly 6. It also defined the size of the arrays, which ranged from 23 kb to larger than 600 kb, surrounded by the same *TP63* flanking sequences identified by LRS. These structures likely arose through sequential rearrangement, potentially driven by homologous recombination, leading to expansion or contraction of tandem units (**Fig. 2D**). Several assemblies (e.g., Assemblies 2, 3, and 4) displayed even more complex structures, revealing multiple tandem arrays interspersed with human genomic segments.

Importantly, OGM also identified a recurrent human-human DNA junction estimated at chr3:189,978,544 and chr3:189,851,767 (hg38, marked by vertical black bars in **Fig. 5B**), which appeared multiple times across different assemblies. This rearrangement closely matches a breakpoint identified in WGS by Amplicon Architect at positions chr3:189,979,893 and chr3:189,849,673 (panel 2, **Fig. 4B**). The minor positional differences likely reflect the spacing between actual breakpoints and adjacent DLE-1 labeling sites, supporting the conclusion that both methods detected the same genomic junction. This junction created a repeat of the 126,777 bp segment that is directly upstream of the HPV-human tandem array. The recurrence of this junction in multiple assemblies, including repeated instances within individual assemblies (Assemblies 2 and 3), strongly suggested origination from a single genomic rearrangement event that then underwent subsequent amplification and restructuring at the megabase-scale.

In summary, OGM revealed two major classes of large-scale DNA structural variation: (1) variation in the number of heterocatemers within tandem arrays, and (2) variation in the number and organization of the larger chromosomal segments containing those arrays. The consistent presence of the same HPV-human junctions, the 7.5 kb viral unit structure, the 2.3 kb deletion human-human junction, and the additional chromosome 3 human-human junction point to a common molecular origin for each distinct rearrangement followed by amplification processes that together reflect extensive genomic instability. Whether all observed assemblies coexist in single SCC47 cells or represent subclonal variation, and whether they are linear integrations within chromosome 3, circular ecDNAs, or a combination of both, remained unresolved.

### FISH analysis reveals mainly chromosomally-inserted, HPV-human heterocatemer DNA plus HPV ecDNA linked with limited or absent human DNA

Fluorescent *in situ* hybridization (FISH) was utilized to determine whether the DNA structures identified by LRS and OGM were intrachromosomal, extrachromosomal, or both. FISH also enabled single-cell analysis of structural variation. Cells were prepared under hypotonic or isotonic conditions. Hypotonic treatment disrupts nuclear membranes, causing metaphase chromosomes to spread, which allows definitive identification of ecDNA. Interphase cells from hypotonic preparations were also analyzed. In contrast, isotonic preparation preserves nuclear integrity and minimizes dispersal of ecDNA. Custom probes targeting HPV16, *TP63,* and *TERC,* which maps approximately 20 Mb upstream of *TP63* on chromosome 3, were generated (**Fig. 6A** and **Supplementary Fig. 4**). Colocalization of HPV16 and *TP63* signals with *TERC* indicated intrachromosomal integration.

**Figure 6.**
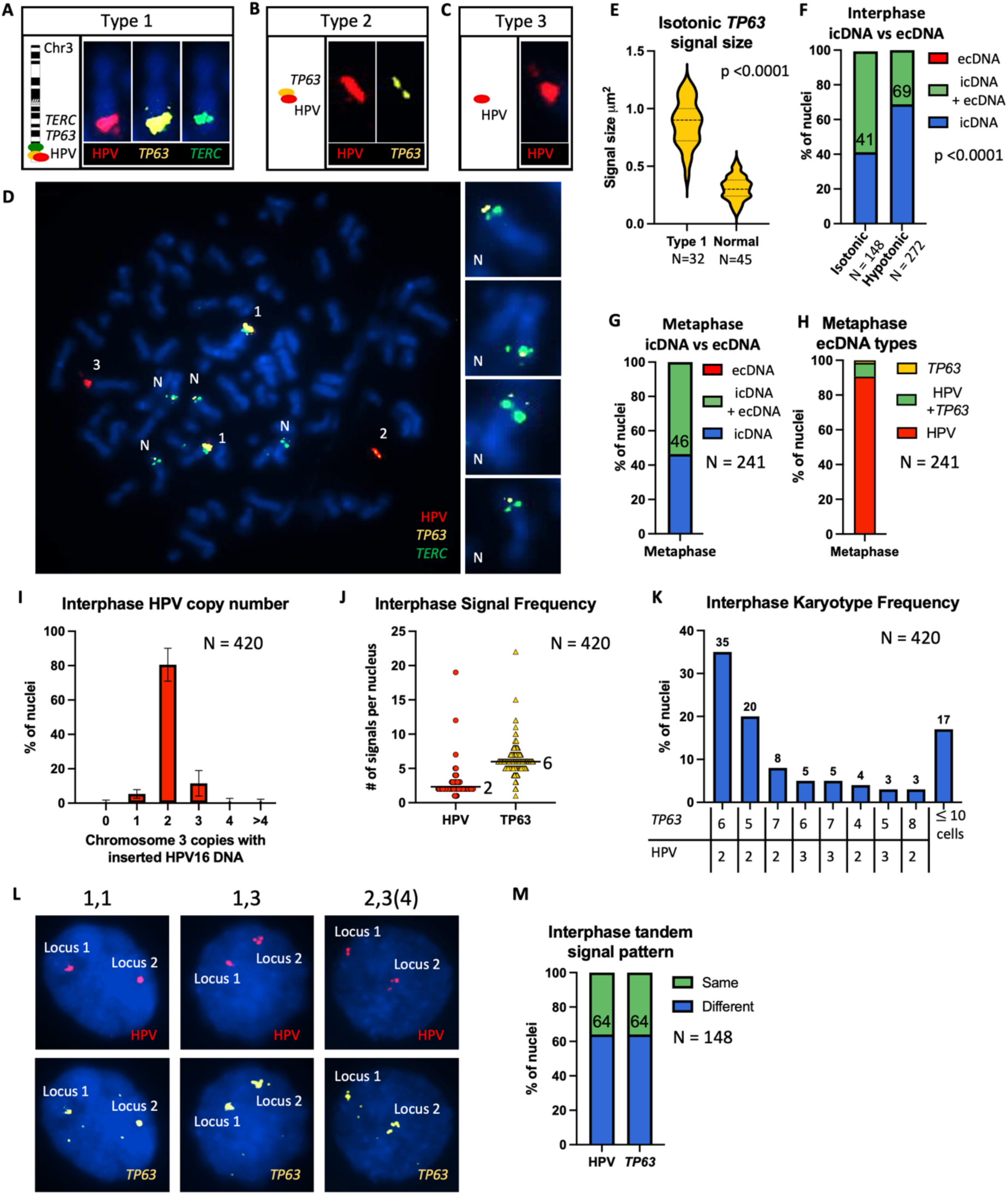
Detection and characterization of intrachromosomal and extrachromosomal HPV16 DNA in single nuclei using FISH: **A)** Type 1 structure: intrachromosomal HPV16 integration in chromosome 3, with colocalization of HPV16 (red), *TP63* (gold), and *TERC* (green) probes. Ideogram shows probe locations on chromosome 3: *TERC* at 3q26.2, *TP63* at 3q28. **B)** Type 2 structure: ecDNA with both HPV16 and *TP63* signals, lacking *TERC.* **C)** Type 3 structure: HPV16-only ecDNA, lacking both *TP63* and *TERC* signals. **D)** Metaphase spread image with merged FISH signals showing all three HPV16 signal types (1–3). Normal chromosome 3 copies without HPV16 but with *TP63* and *TERC* are labelled “N”. **E)** Quantification of *TP63* FISH signal size between Type 1 loci (N= 32) and normal chromosome 3 loci (N=45) in interphase nuclei. *TP63* signals at Type 1 loci are significantly larger (T-test, *p*<0.0001). **F)** Frequency of intrachromosomal (icDNA) and extrachromosomal (ecDNA) HPV16 signals in isotonic (N = 148) and hypotonic (N = 272) interphase nuclei. Blue: icDNA only; green: both icDNA and ecDNA; red (not detected): ecDNA only. icDNA-only frequencies are shown above the blue bars. A significant difference was observed between isotonic and hypotonic preparations (Fisher’s exact test, p<0.0001). **G)** Frequency of icDNA and/or ecDNA in metaphase nuclei. **H)** Classification of metaphase nuclei based on ecDNA content: HPV16 only (red), HPV16 plus *TP63* (green), or *TP63* signal only (*gold*). N = 241. **I)** Distribution of the number of copies of chromosome 3 with intrachromosomal HPV16 DNA per interphase nucleus (pooled isotonic and hypotonic). N = 420. **J)** Total number of HPV16 (red) and *TP63* (gold) FISH signals per interphase nucleus. N = 420. **K)** Karyotype distribution showing number of chromosome 3 copies with and without HPV16 DNA. Bars show percentages and are annotated with the corresponding number of HPV16 and *TP63* signals. Karyotypes found in fewer than 10 nuclei are grouped in the final column. N = 420. **L)** Representative interphase nuclei showing different tandem signal patterns at two HPV16 integration loci. Patterns are labelled based on number of punctate signals per locus (e.g, 1,1 or 3,4). **M)** Frequency of hypotonic interphase nuclei (N = 148) with differing (blue) versus identical (green) numbers of HPV16-*TP63* tandem signals between two integration loci. Percentage of nuclei with differing signals is shown above each blue bar.

Three distinct types of HPV16 signals were observed among SCC47 metaphase spreads. Type 1: HPV16, *TP63,* and *TERC* signals, consistent with HPV16 intrachromosomal integration (**Fig. 6A**). Type 2: HPV16 and *TP63* without *TERC,* suggesting HPV1-*TP63* ecDNA (**Fig. 6B**). Type 3: HPV16 alone, representing ecDNA with undetectable *TP63* signal (**Fig. 6C**). HPV16-negative copies of chromosome 3 containing *TP63* and *TERC* signals were also detected, representing normal copies of the chromosome. An example of a metaphase spread containing all three types of HPV16 signals and normal copies of chromosome 3 is shown in **Fig. 6D**. *TP63* signals at integrated loci (Type 1) were significantly larger than those observed at non-integrated loci (N in **Fig. 6D**) (t-test of isotonic interphase nuclei, *p*<0.0001, **Fig. 6E**). This suggested that chromosomally-integrated loci harbor the tandemly repeated HPV16-*TP63* DNA including the 16 kb *TP63* exon 12 through 16 segment detected by OGM and LRS, whereas the HPV16-negative copies of chromosome 3 copies retain a single, unamplified *TP63* locus.

In 100% of 420 interphase nuclei (148 isotonic and 272 hypotonic), HPV16-*TP63* heterocatemer FISH signals colocalized with chromosome 3 (**Fig. 6F**), unambiguously demonstrating intrachromosomal integration as the dominant form. Metaphase spreads also showed 100% of nuclei with intrachromosomal integration (**Fig. 6G**). No nucleus contained only HPV16-*TP63* ecDNA, although approximately half of the nuclei contained both intrachromosomal DNA and ecDNA (green bars, **Fig. 6F-G**), with higher ecDNA detection under isotonic conditions (Fisher’s exact test, *p* <0.0001), likely due to reduced dispersal of ecDNA under those conditions.

Most ecDNA signals contained HPV16 without detectable *TP63*, fewer contained both HPV16 and *TP63*, and a minority showed *TP63* alone (**Fig. 6H**). The absence of *TP63* in most ecDNAs may indicate true absence or signal below FISH detection limit. In either case, these structures likely do not correspond to the large tandem HPV16-*TP63* heterocatemers observed by LRS and OGM (**Figs. 2** and **5**). Because the 406 bp HPV16 E2 segment was absent from all HC+SEQ, LRS, and WGS reads, the ecDNAs detected by FISH likely originated from the rearranged intrachromosomal form lacking the E2 segment (**Fig. 1**). We hypothesize that these ecDNAs arose from DNA replication intermediates on intrachromosomal templates that circularized at random positions (**Fig. 2D**).

Variability in chromosome 3 copy number and integration status was also observed. Most nuclei (79 ± 9%) had two copies with intrachromosomal HPV16 integration (**Fig. 6I**). Additionally, all nuclei retained copies without an integration, with a median of six total *TP63* signals per nucleus (**Fig. 6J**). The most common karyotype (35%) consisted of two integrated and four non-integrated copies (**Fig. 6K**). Over 60% of nuclei had two copies of chromosome 3 with an HPV16 integration, plus an additional three to five copies lacking HPV16 but retaining presumably intact *TP63*. Interestingly, chromosome 3 counts ranged from a minimum of 4 to more than 10 copies per nucleus (**Supplementary Fig. 6B**), reflecting extensive karyotypic heterogeneity.

Under isotonic conditions, intrachromosomal DNA loci often contained multiple linear FISH signal repeats for HPV16 and *TP63*, consistent with linked heterocatemer tandem arrays such as those detected by OGM. Quantification of the punctate signals allowed the assessment of tandem unit counts within each nucleus (**Fig. 6L**). Since almost all nuclei had at least two integrated chromosome 3 copies, we compared tandem counts within individual nuclei. In most cases, one locus contained more tandem signals than the other (**Fig. 6M** and **Supplementary Fig. 6**, Wilcoxon signed ranks, p<0.0001), revealing intra-nuclear, tandem copy number heterogeneity at the *TP63* locus. This asymmetry could indicate that one chromosome 3 copy with HPV integration was duplicated and subsequently underwent independent structural evolution, including ongoing recombination, unequal amplification/contraction, or additional types of genomic instability.

The large FISH signals (**Fig. 6D and E**) are interpreted to be multiple HPV-*TP63* copies in a compact structure that precludes identification of individual segments. To test this, we treated SCC47 with the bromodomain and extra-terminal (BET) protein inhibitor JQ1. BET proteins perform key roles in nuclear structure and transcription, and JQ1 is known to inhibit HPV-transformed cells(63,64). Following treatment, we observed a reproducible increase in the spatial separation and elongation of HPV16 FISH signals at the *TP63* locus, consistent with decompaction or relaxation of the structure of the integration loci (**Fig. 7A**). Quantification of JQ1-treated versus control nuclei showed a significant increase in stretched or linearly extended FISH signal patterns (**Fig. 7B**). These results suggest that the compact topological structures of the HPV-*TP63* DNAs within chromosome 3 depend on BET proteins.

**Figure 7.**
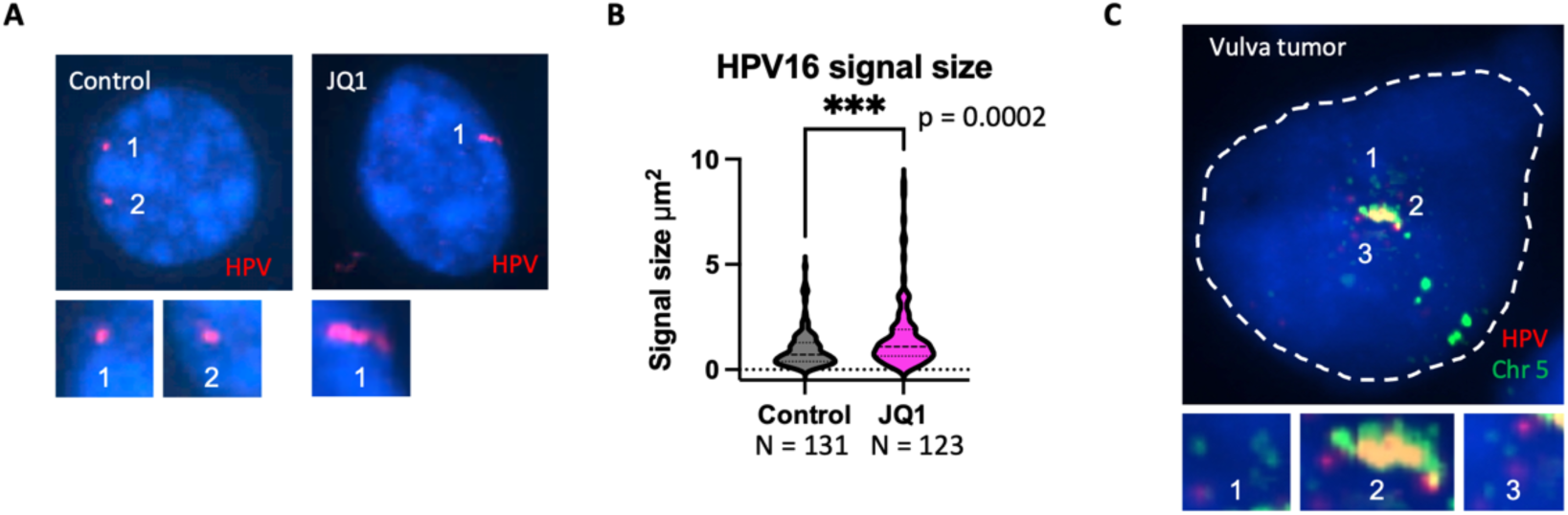
BET inhibitor effect on integrated HPV16 signals and comparison to tumor tissue FISH: **A)** Representative interphase FISH images of control versus cells treated with 1μm JQ1 showing HPV16 signals in red. **B)** Quantification of HPV16 signal size in control (grey, N=131) versus JQ1 (pink, N=123) cells showing a significant increase in size after treatment (Mann-Whitney U test, p=0.0002). **C)** FISH image of an interphase nucleus from a primary vulvar squamous carcinoma showing HPV16 (red) integrated into chromosome 5 (green). Positions 1 and 3 represent ecDNAs; position 2 shows a large intrachromosomal HPV16 integration.

Finally, SCC47 is a cultured cell line, and our findings raise the question of whether similar changes occur in primary tumors *in vivo*. Therefore, we performed FISH on a primary vulvar cancer harboring an HPV16 integration on chromosome 5. We observed a strong, compact intrachromosomal HPV16 signal together with ecDNA in a fraction of tumor nuclei, an example of which is shown in (**Fig. 7C)**. This pattern mirrors that observed in SCC47 and supports the possibility that these structures occur in primary tumors.

In summary, FISH analysis demonstrated that most HPV16-human DNA structures were intrachromosomally integrated at the *TP63* locus on chromosome 3 and that approximately half of the cells also contained extrachromosomal DNA. Moreover, we observed substantial genomic variability in both copy number and HPV16-*TP63* signal patterns across nuclei. Notably, treatment with the BET bromodomain inhibitor JQ1 led to visible stretching and spatial separation of the HPV16 and *TP63* FISH signals, suggesting that the integration locus resides in a chromatin context that structurally involves BET proteins. Together, these findings reinforce a model in which HPV DNA integration can ultimately produce large, heterogeneous, structurally compact HPV-human DNA structures that vary not only among individual cultured SCC47 cancer cells, but also between homologous chromosomes, and may be relevant to HPV-associated cancers *in vivo*.

## DISCUSSION

This study leveraged a multi-resolution genomic approach to determine the complex architecture of HPV16 DNA integration in a cancer-derived cell line. Combining high-resolution sequencing analyses with custom-designed FISH probes provided an approach to link nucleotide-level structure with chromosomal context, establishing a platform capable of resolving oncogenic DNA architectures and defining HPV-human DNA heterocatemer organization, including novel large-scale rearrangements. In particular, the FISH analysis, perhaps unexpectedly, determined that the HPV-human heterocatemers were intrachromosomal in 100% of the cells. In addition, the inherent spatial localization capability of FISH combined all the HPV16 DNAs in the cell cultures lacked 406 bp of the E2 gene, with the nucleotide-level resolution of sequencing and mapping technologies established a structural relationship between intrachromosomal HPV-human heterocatemers and ecDNAs in the same cells. Thus, this study establishes a broadly applicable strategy for distinguishing ecDNA from intrachromosomal DNA in cancers, which can be challenging to accomplish by sequencing alone.

Previous studies have described viral integration sites, viral-human DNA junctions, and tandem repeat structures (4,9,10,12,13,33), plus the presence of extrachromosomal HPV DNAs in tumors(14,15,65–67). However, short-read sequencing alone cannot distinguish a putative circular DNA from tandemly repeated, chromosomally integrated units with identical sequences. By combining LRS and OGM, we identified 23.5 kb heterocatemers comprising HPV16-*TP63* DNA sequences arranged in large tandem arrays extending up to 0.6 Mb (**Fig. 5B** and Assembly 6, **Supplementary Fig. 3**). These analyses also identified complex genomic rearrangements involving flanking human sequences spanning from hundreds of kilobases to over a megabase, reinforcing the value of long-read genomic technologies for capturing higher-order structural variation. Nevertheless, neither LRS nor OGM definitively determined whether these large, rearranged structures were extrachromosomal or intrachromosomal.

FISH analysis unambiguously demonstrated that the large HPV16-*TP63* tandem arrays were chromosomally integrated at the *TP63* locus on chromosome 3 in all nuclei analyzed. Concurrently, FISH detected HPV16-containing ecDNA in over half of the cells, predominantly consisting of HPV16 sequences with no detectable *TP63*, although a subset also contained *TP63* DNA. In addition, FISH revealed considerable chromosome 3 copy number variation and intranuclear heterogeneity in HPV16 and *TP63* signal patterns (**Fig. 6** and **Supplementary Fig. 6**). Given that *TP63* is frequently amplified in HPV-related cervical cancers(68–70), even in the absence of integration at this locus, the presence of multiple chromosome 3 copies with unaltered *TP63* may preserve essential functions of *TP63* in transformed keratinocytes(62,69,71). Importantly, the marked variability in signal intensity, copy number, and composition across different copies of chromosome 3 within nuclei highlights the need for spatially-resolved genomic methods like FISH to capture the full spectrum of DNA architecture and subclonality in cancer.

The wide range of HPV16-human DNA structures observed can be mechanistically explained by known models of viral integration and amplification. Holmes et al.(54) proposed that HPV genomes whose ends are oriented toward each other may initially integrate through a non-linear mechanism, forming circular intermediates. Our data support this model and extend it to explain how downstream complex integration and amplification patterns may form. In SCC47, the HPV16 ends joined to *TP63* are inwardly oriented (**Fig. 1**), consistent with formation of an extrachromosomal circular HPV16-*TP63* intermediate. We propose that this circular intermediate subsequently underwent homologous recombination with the unaffected *TP63* locus on chromosome 3, leading to an integrated structure with flanking direct repeats of *TP63* (**Fig. 2D**). Subsequent rounds of amplification likely occurred via DNA polymerase slippage or non-allelic homologous recombination between the flanking human repeats(72–75), yielding the large tandem arrays observed. Whenever full tandem arrays were recovered in long-read sequences, the human flanking regions were identical on both ends, supporting this mechanism (**Fig. 2** and **Supplementary Fig. 2**). OGM further revealed copy number variation within arrays, consistent with dynamic recombination-based expansion and contraction.

FISH analysis further revealed variable DNA content within ecDNA, including HPV16-only signals, HPV16 colocalized with *TP63*, and *TP63*-only signals (**Fig. 6** and **Supplementary Fig. 5**). While FISH cannot always distinguish true absence from signals below detection threshold, the consistent appearance of distinct patterns across nuclei, and their absence in others, strongly supports the existence of actual molecular heterogeneity rather than low-level signal loss. Given the ultra-deep coverage of HC+SEQ, far greater than WGS (**Fig. 1A**), and the fact that all the HPV16 DNAs in the cell cultures lacked 406 bp of the E2 gene, we infer that the ecDNAs arose, at least in part, from the chromosomally integrated structures. Incompletely replicated HPV DNAs have been proposed as intermediates in HPV DNA integration events(28), and we speculate that these could similarly serve as substrates for circularization into ecDNAs (**Fig. 2D**). Circularization at different positions within incompletely replicated HPV16-*TP63* heterocatemer DNAs could account for the heterogeneity observed by FISH. Alternatively, homologous recombination between flanking human repeats, or indeed any repeated sequences, might lead to excision of heterocatemer units from intrachromosomal tandem units, thus generating ecDNAs. These novel data establish a relationship between ecDNA and intrachromosomal DNA and may provide a mechanism that could contribute to maintenance of ecDNA regardless of how efficiently ecDNA independently replicates and segregates to daughter cells.

Once formed from intrachromosomal DNA, a key unresolved question is how the ecDNAs are maintained in SCC47 cells. One possibility is that host-mediated mechanisms, potentially similar to those used by oncogenic human-derived ecDNA(23,66,76), might contribute to the maintenance of HPV-*TP63* ecDNA in SCC47. A second possibility is that HPV-specific processes contribute to ecDNA maintenance. During productive HPV infection, high-copy-number replication of circular HPV DNA relies on the viral E1 and E2 proteins and a replication initiation site in the URR(77,78). However, most HPV tumors, including SCC47, lack an intact E2 ORF, presumably precluding HPV-mediated replication. A third speculative possibility is that ecDNAs are constantly regenerated from intrachromosomal DNAs, a possibility consistent with the heterogeneity of the ecDNA observed here by FISH. Notably, circular DNA arising from intrachromosomal tandem repeats has been documented in non-HPV tumors(79), and a similar mechanism might contribute to the maintenance of extrachromosomal DNAs in SCC47 cells. How much each or all of these are involved in ecDNA maintenance deserves further study. Regardless of the mechanisms involved, the persistence of the HPV-containing ecDNAs could contribute to amplification of the HPV16 E6 and E7 oncogenes in a significant fraction of cells, enhancing their tumorigenic potential.

RNA transcripts containing the E6 and E7 ORFs in SCC47, particularly those detected by long-read RNA-seq, could plausibly be transcribed from either intrachromosomal or extrachromosomal templates (**Fig. 3**). The HPV-human chimeric transcripts that were polyadenylated at the *TP63* exon 16 site could only derive from templates at the 3’ end of a tandem array that include the *TP63* sequence downstream of the heterocatemer, such as the intrachromosomal, tandem heterocatemer structures (**Fig. 3B**). Moreover, our long-read RNA-seq was also consistent with some transcripts originating within an internal tandem unit (**Fig. 3B**). This strongly argues that internal units within the tandem arrays can be transcriptionally active.

Extrachromosomal DNAs are widespread in human cancers and have been associated with poor clinical outcomes and resistance to antitumor therapies(23,24,66). Oncogenes are frequently present on ecDNAs(24,80), and in HPV-driven cancers, the E7 and E6 oncogenes can be within the viral segments that form these circular structures(4,5). The finding of intrachromosomal and ecDNAs is also reminiscent of double-minute (DM) extrachromosomal and homogeneous staining region (HSR) intrachromosomal DNAs, both of which are associated with resistance to DNA-damaging cytotoxic therapies (e.g., cisplatin, doxorubicin) as well as reduced responses to immuno-oncology agents (e.g., anti-PD-1 checkpoint inhibitors such as nivolumab or pembrolizumab) (81–83). In SCC47 cells, we observed primarily intrachromosomal tandemly repeated HPV16-human heterocatemers with a subset of cells also containing ecDNA structures. We suggest that the coexistence of both forms and the heterogeneous distribution of ecDNA in SCC47 may support phenotypic diversity, adaptation, and possibly evolution of subclones.

Although this study focused on a cultured cell line derived from an HPV-induced tumor, whether similar DNA structures exist in primary HPV-induced tumors remains an open question. However, our analysis of a primary vulvar squamous cell carcinoma(4) (and **Fig. 7C**) revealed strong intrachromosomal HPV16 FISH signals on human chromosome 5 coexisting with extrachromosomal signals, suggesting that this phenomenon may extend beyond cell lines. Our results highlight the need for caution when interpreting sequencing data, as ecDNA and intrachromosomal tandem repeat structures can yield indistinguishable readouts in many genomic assays. Additional techniques, such as FISH (**Fig. 6**)(24,25), are essential for determining DNA topology and for identifying cases in which both extrachromosomal and intrachromosomal forms coexist. The methodology used here to reveal extensive structural variation associated with HPV DNA integration in cancer genomes could be applied more broadly to investigate multi-scale DNA structures in other cancers, including determining whether highly amplified, HPV and/or human-derived oncogenes share common organizational principles.

## ACKNOWLEDGEMENTS

We thank Adriana Messyasz from the Rutgers NJMS Molecular and Genomics Informatics Core for assisting with bioinformatic analysis of LRS assemblies. Services (RNA extraction), results, and products in support of the research project were generated by the Rutgers Cancer Institute, ImmuneMonitoring, and Flow Cytometry Shared Resource, supported, in part, with funding from the NCI-CCSG P30CA072770-5920. Services, results, and/or products supporting this research were provided by the Biospecimen Repository and Histopathology Service Shared Resource at the Rutgers Cancer Institute of New Jersey, supported in part by funding from NCI-CCSG P30CA072720-25. The authors acknowledge the Office of Advanced Research Computing (OARC) at Rutgers, The State University of New Jersey for providing access to the Amarel cluster and associated research computing resources that have contributed to the results reported here.

## AUTHOR CONTRIBUTIONS

**Eleanor J. Agosta**: Investigation; Formal analysis; Visualization; Writing – original draft. **Yoke-Chen Chang, Vandya Rao, Jessie Hollingsworth, Michelle Brown**: Investigation; Data curation; Formal analysis. **Debajyoti Kabiraj, Chang Chan, Danny E. Miller, Brian J. Haas**: Formal analysis; Software; Data curation. **Brian J. Haas**: Methodology. **Advaitha Madireddy, Subhajyoti De, Mark Einstein, Anne Van Arsdale, Koenraad Van Doorslaer**: Conceptualization; Data interpretation; Writing – review & editing. **Koenraad Van Doorslaer**: Resources. **Jack Lenz, Cristina Montagna**: Conceptualization; Funding acquisition; Project administration; Supervision; Writing – review & editing; Visualization.

## SUPPLEMENTARY DATA

Supplementary Data are available at NAR online.

## DATA AVAILABILITY

Sequencing data generated in this study are available in the SRA of the National Library of Medicine, National Institutes of Health, under the accession number PRJNA1247532 and are publicly available as of the date of publication. The dataset includes FASTQ files obtained from: targeted enrichment of HPV DNA obtained from custom hybridization capture probes to HPV16; whole genome pair-end sequencing; RNA sequencing; and long-read DNA and RNA sequencing performed using the Oxford Nanopore library prep kits SQK-LSK110 and SQK-PCS110 following manufacturer instructions. Optical genome mapping data have been deposited at Zenodo and are publicly available as of the date of publication at 10.5281/zenodo.16781564. This paper does not report original code. Any additional information required to reanalyze the data reported in this paper is available by the lead contact, Cristina Montagna, upon request.

