## Supplemental for "Complex HPV-human DNA structures revealed by large-scale DNA analyses in an HPV-cancer derived cell line"

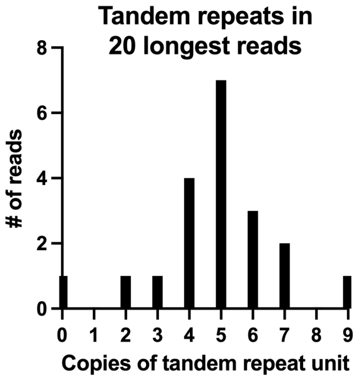


**Supplementary Figure 1. Number of tandem repeat units in the 20 longest, nanopore sequence reads.** For the 20 longest reads that contained HPV16 DNA, the number of sequence reads is plotted as a function of the number of tandem units shown on the X-axis. N=20.

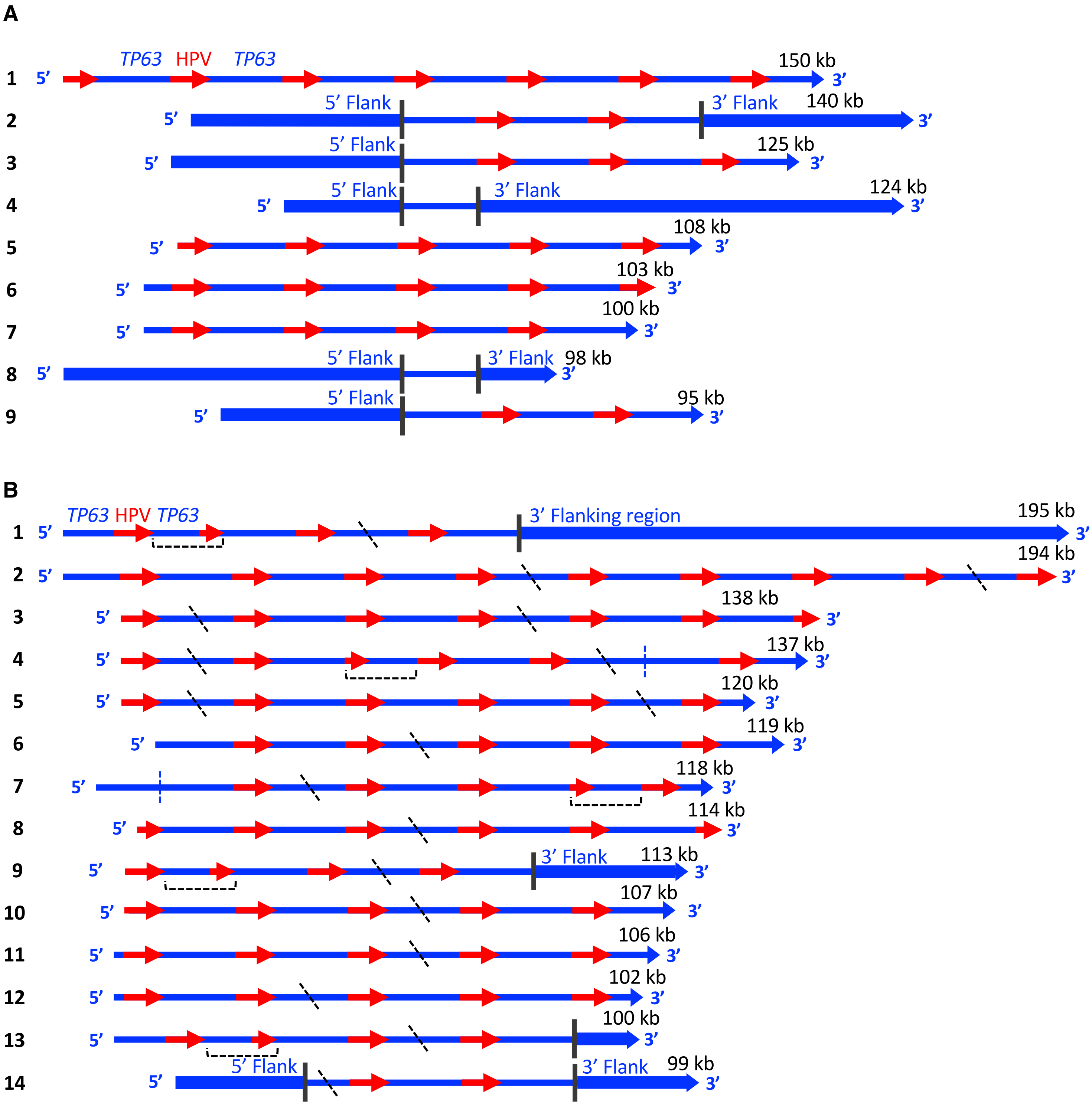


**Supplementary Figure 2. Maps of the 20 longest nanopore reads at the *TP63* locus in SCC47 cells. A)** Maps of reads without gaps or truncations are shown. The length of each individual sequence read is shown. HPV16 segments are in red and human segments are in blue. 5' and 3' flanking regions that map upstream and downstream of *TP63* exons 12-16 are indicated by thick blue segments and vertical black lines. **B)** Maps of reads containing gaps or truncations. HPV16 segments are in red and human segments are in blue. 5' and 3' flanking regions are indicated by vertical black lines and thick blue segments. Diagonal dotted black lines represent ~2.3 kb gaps defined in Fig. 2E. Brackets underneath reads represent truncations of HPV16 and *TP63* segments in individual reads. The vertical blue dotted lines represent human-human junctions.

A

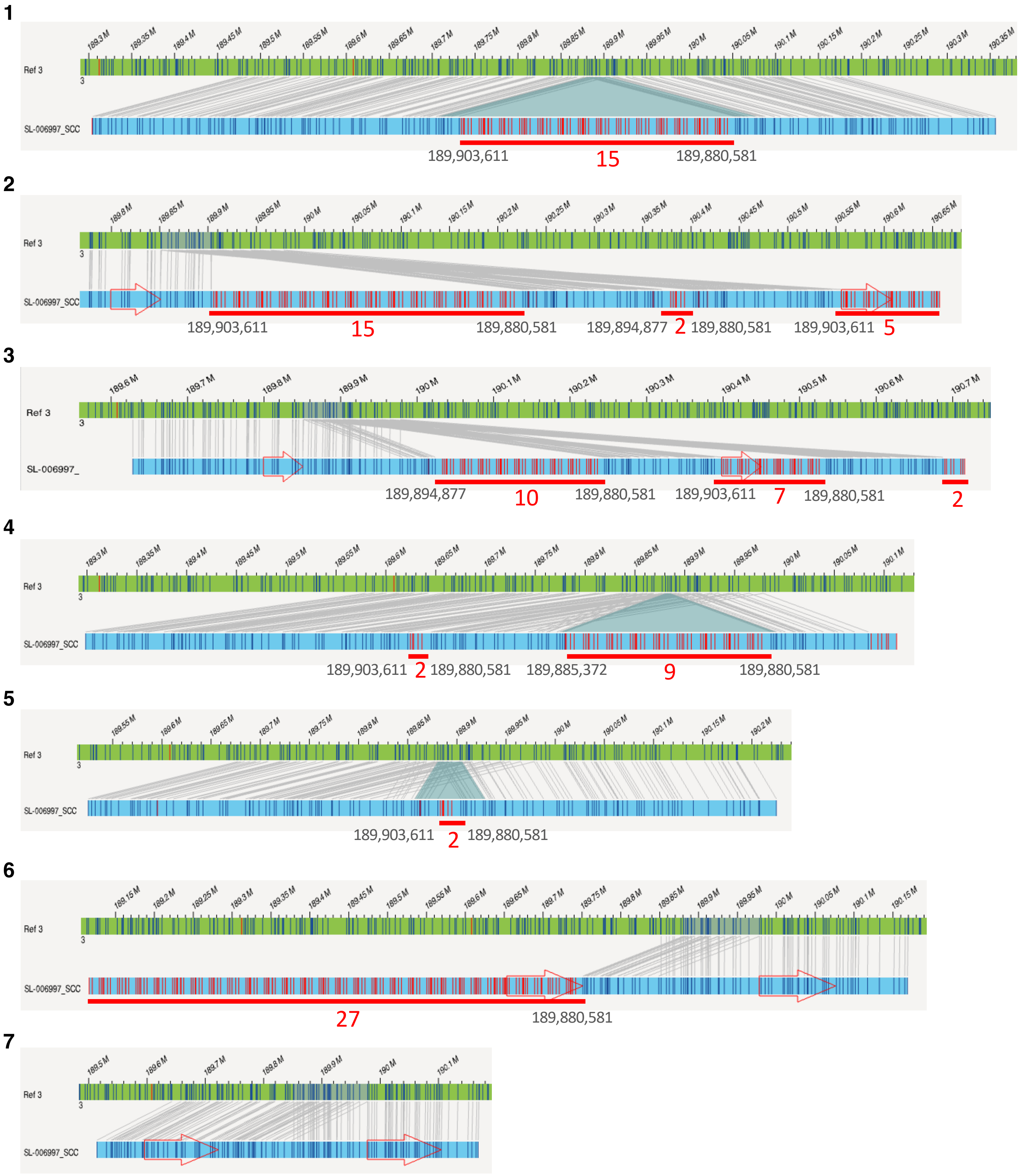


**Supplementary Figure 3. Detailed maps of each of the 7 the optical genome mapping assemblies.** Chromosome 3 is shown in green with genomic locations shown above and DLE1 labeling sites represented by dark blue vertical lines. Each assembly is shown in light blue with dark blue lines showing DLE1 sites that align to chromosome 3. Gray lines align the assemblies with chromosome 3. Vertical red lines showing DLE1 sites from exons 12-16 of *TP63* on chromosome 3. Each tandem array is marked by a horizontal red line with the number of tandem HPV16-*TP63* units shown underneath. The genomic location of the DLE 1 sites at the beginning and end of each tandem array are shown underneath in gray. The large red arrows on various assemblies indicate the direction of ascending genomic positions from hg38.

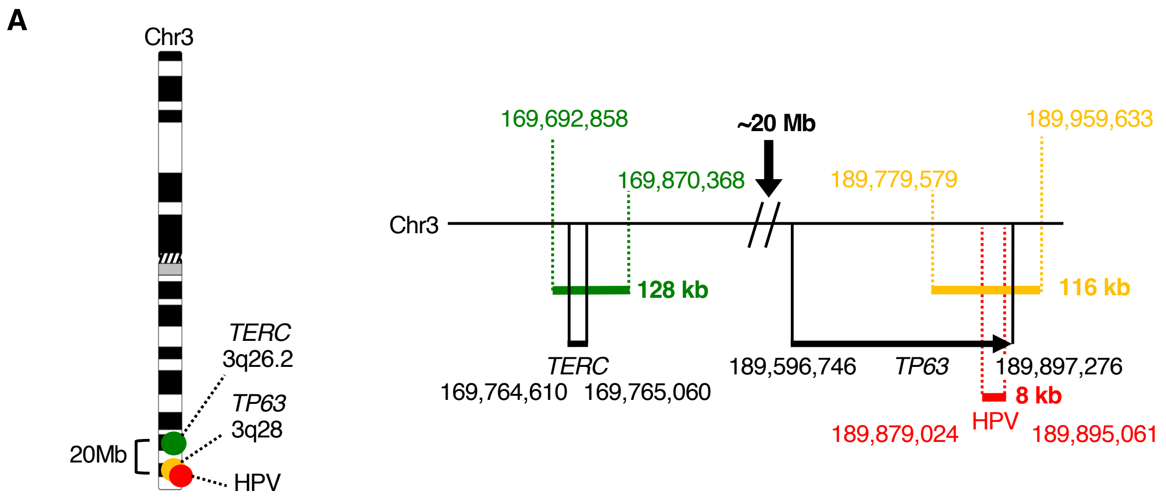


**Supplementary Figure 4. Location of BAC clones used to make each custom FISH probe.** Ideogram view shows an overview of the location of each of the probes with *TERC* in green, *TP63* in gold, and HPV16 in red. Detailed map on the right shows the specific location of each BAC clone used to make custom probes. Thick colored lines represent the probe location with sizes to the left. Specific genomic locations on chromosome 3 indicated by dotted lines and labeled with genomic coordinates. Solid black lines represent the *TERC* and *TP63* gene locations with specific coordinates on either side of the gene name. The HPV16 DNA segment is positioned below exons 12-16 of *TP63*. The 20 Mb space between *TERC* and *TP63* is indicated.

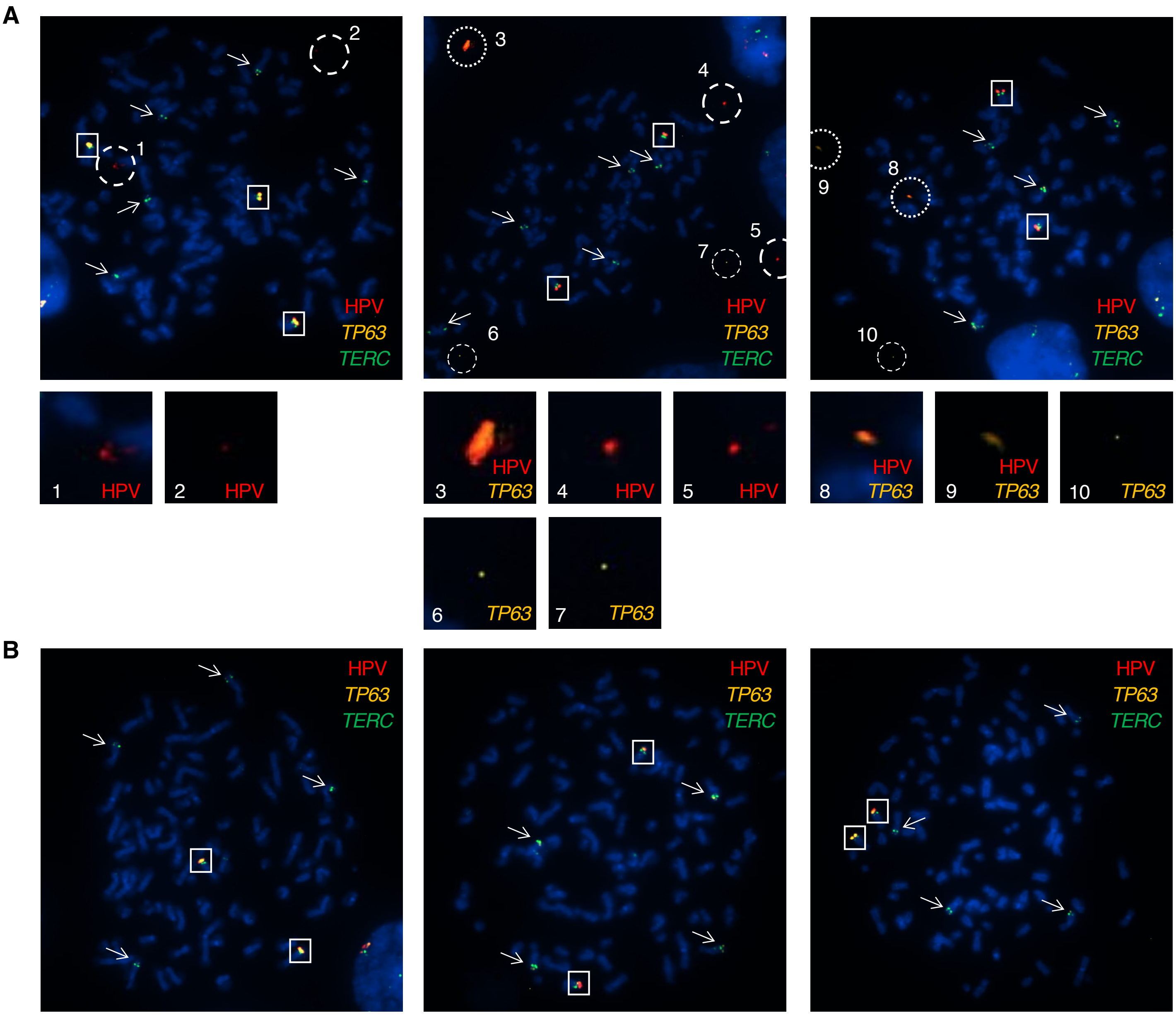


**Supplementary Figure 5. Representative metaphase spread images with and without extrachromosomal DNA. A)** Representative images of three metaphase nuclei with extrachromosomal DNA. Dashed large circles represent HPV-only signals. Dotted large circles represent overlapping HPV and *TP63* signals. Small dashed circles representing *TP63* only signals. Boxes represent copies of chromosome 3 that contain intrachromosomal HPV signals and arrows represent copies of chromosome 3 without HPV signals. The small panels represent enlarged images of the indicated, numbered loci. **B)** Representative images of three metaphase nuclei without extrachromosomal DNA. Boxes represent copies of chromosome 3 that contain HPV signals, and arrows represent copies of chromosome 3 without HPV signals.

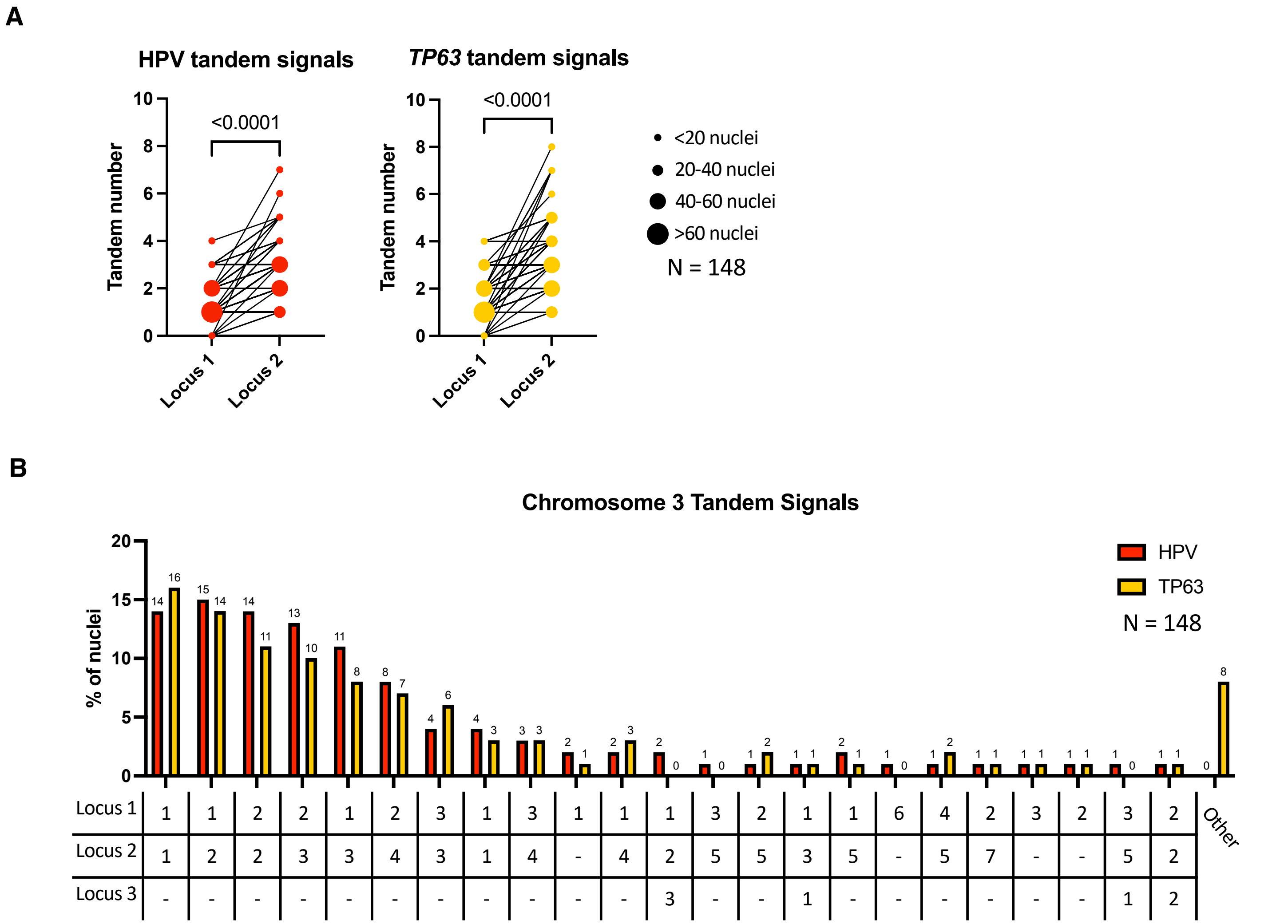


**Supplementary Figure 6. Comparison of tandem signal number on each HPV16 DNA-containing copy of chromosome 3 in an individual nucleus. A)** Tandem signals of HPV or *TP63* between two loci in interphase cells as paired data points. Bubble size represents the number of loci sharing that number of tandem signals.Lines connecting bubbles represent specific pairwise combinations. Analysis was done on hypotonic interphase nuclei N =148. Cells that had more than two loci with HPV integrations had only the two loci that differed from each other included. Comparison of the numbers of signals between locus 1 and locus 2 showed a significantly higher number of signals at locus 2 for each probe ($p < 0.0001$, nonparametric Wilcoxon matched-pairs signed rank test). **B)** Frequency of specific tandem signal patterns at each chromosome 3 locus in individual nuclei. HPV signal data are plotted in red and *TP63* signal data are in gold. The number of signals at each locus is represented in the table below each pair of bars. The larger number was always arbitrarily denoted as locus 2. A few nuclei had three HPV16-containing copies of chromosome 3 which were denoted as locus 3. The percentage of nuclei with each pattern is specified in black numbers above each bar. The remaining signal patterns that occurred in 1% or fewer nuclei are grouped into the “other” category. Analysis done on hypotonic interphase nuclei N =148.

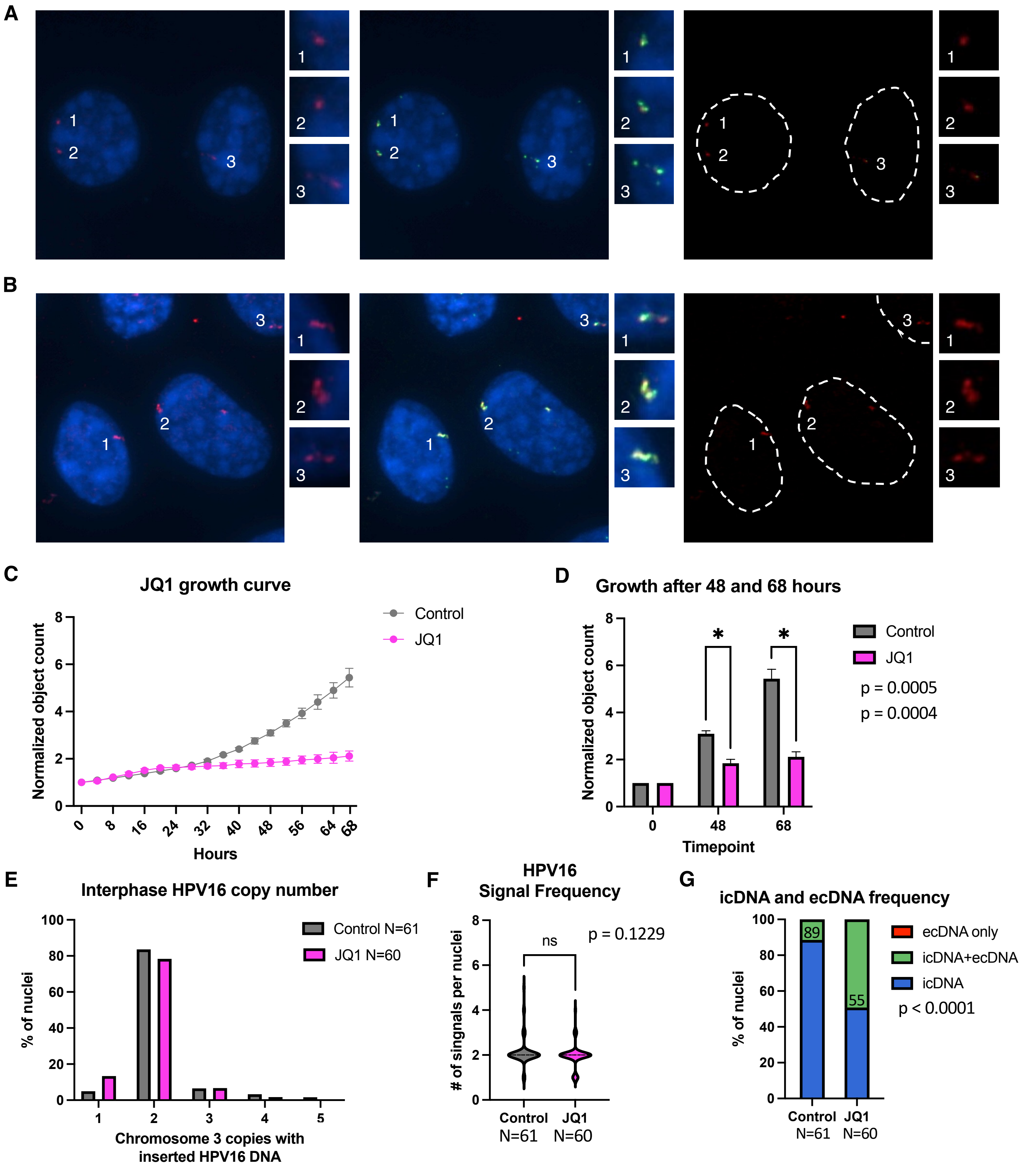


**Supplementary Figure 7. SCC47 cells treated with JQI images and quantification A)** Representative images showing HPV (red) and *TP63* (gold) signals of untreated cells **B)** Representative images of cells treated with 1 µm JQ1. **C)** Growth curve of control cells (grey, N=4 wells) and treated with 1 µm JQ1 (pink, N=2 wells). **D)** Normalized object count between control (grey) and 1 µm JQ1 cells (pink) at 48 and 68 hours (p=0.0005 and p=0.0004, student's t-test). **E)** Distribution of chromosome 3 copies with an inserted HPV16 signal by FISH. Control in grey (N=61 cells), 1 µm JQI treatment (N=60) in pink. **F)** Total number of HPV16 FISH signals per control (grey, N=61) and JQI (pink, N=61) treated cells. (HPV16 p=0.1229, nonparametric Wilcoxon matched-pairs signed rank test). **D)** Frequency of intrachromosomal (icDNA) and extrachromosomal (ecDNA) HPV16 signals in Control (N = 61) and 1 µm JQI (N = 60) nuclei. Blue: icDNA only; green: both icDNA and ecDNA; red (not detected): ecDNA only. icDNA-only frequencies are shown above the blue bars. A significant difference was observed between control and JQI cells (Fisher's exact test, p<0.0001).
